# Multiplexed measurement of protein biomarkers in high-frequency longitudinal dried blood spot (DBS) samples: characterization of inflammatory responses

**DOI:** 10.1101/643239

**Authors:** N. Leigh Anderson, Morteza Razavi, Matthew E. Pope, Richard Yip, Terry W. Pearson

**Affiliations:** SISCAPA Assay Technologies, Inc. PO Box 53309, Washington DC 20009

## Abstract

A detailed understanding of changes in blood protein biomarkers occuring in individuals over time would enable truly personalized approaches to health and disease monitoring. Such measurements could reveal smaller, earlier departures from normal baseline levels of biomarkers thus allowing better disease detection and treatment monitoring. Current practice, however, generally involves infrequent, sporadic biomarker testing, and this undersampling likely fails to capture important biological phenomena. Here we report the use of a robust multiplex immuno-mass spectrometric method (SISCAPA) to measure a panel of clinically-relevant proteins in a unique collection of 1,522 dried blood spots collected longitudinally by 8 individuals over periods of up to 9 years, with daily sampling during some intervals. Analytical workflow CVs of 2-6% for most assays were achieved by normalizing DBS plasma volume using a set of 3 minimally varying proteins, facilitating temporal analysis of both high- and low-amplitude biomarker changes compared to personalized baselines. The biomarkers included a panel of 9 positive and 5 negative acute phase response (inflammatory) proteins, allowing longitudinal analysis of inflammation markers associated with major and minor infections, influenza vaccination, recovery from hip-replacement surgery and Crohn’s disease. The results illustrate complex time-dependent “biomarker trajectories” on multiple timescales and provide a basis for detailed personalized models of inflammation dynamics. The striking stability of most biomarker protein levels over time, combined with the convenience of self-sampling and low cost of multiplexed measurements using mass spectrometry, provide a new window into the temporal dynamics of disease processes. The extensive results obtained using this high throughput approach offer a new source of precision biomarker ‘big data’ amenable to machine learning approaches and application to more personalized health monitoring.

## Introduction

Most important events in human health unfold as quantitative changes that occur on timescales of hours, days, months and years. Modeling, classifying and predicting such changes in individuals is a central goal of personalized medicine, and requires access to longitudinal measurements of important molecular parameters (i.e., biomarkers) before, during and after health- and disease-related events. Unfortunately, current biomarker approaches grossly undersample these dynamic events, providing only a low-resolution picture of health and disease at the molecular level. While advances in wearable sensors [1], coupled with developments in machine learning [2], have triggered an explosion of interest in the potential of “big data” to shed light on health problems [3], the application of this approach to track events at the molecular level has been limited by the scarcity of dynamic readouts of the body’s internal molecular sensors, represented, for example, by the levels of biomarker molecules (principally proteins) in the blood. Although more than 100 clinical laboratory tests are available to measure specific blood protein biomarkers [4], such tests are typically used one at a time and only when an immediate medical diagnostic need justifies the cost. As a result, there is a paucity of protein biomarker data measured in individuals at frequencies matching the dynamics of health events, and even less data using panels of biomarkers that cover multiple aspects of a person’s physiology.

To fill this gap we sought to develop a practical approach to obtain precise measurements of blood biomarkers at high frequency and low cost, using samples that can easily be collected by subjects themselves at any time and any place: dried blood microsamples. In the work described here we demonstrate the use of such a method measuring pre-selected health-relevant proteins by SISCAPA immuno-mass spectrometry [5] in conventional dried blood spot (DBS) samples collected on filter paper cards over spans of years, including extensive periods of daily collection. While this approach could be considered excessive in the context of most routine diagnostic applications, it has enormous potential value when applied to specific periods of time during which significant health events are anticipated: e.g., disease progression studies, clinical trials, surgery, or pre-arranged changes in medication, lifestyle or diet [6].

Dried whole blood samples, in the form of DBS on filter paper cards, have been used as clinical samples since the pioneering studies of Guthrie [7] on inborn errors of metabolism, and have slowly gained acceptance for a few narrow applications in epidemiological studies, pharmaceutical trials and direct-to-consumer testing. However, the widespread use of DBS as an alternative to conventional phlebotomy has been limited by the difficulty of determining the serum- or plasma-equivalent volume present in a given area of filter paper (typically a circular 6 mm diameter punch) excised from the dried blood spot. The blood volume in a standard filter paper punch can vary by +/-10-15%, while the fraction of this blood consisting of plasma can also vary significantly (∼30-60%) as a function of the sample hematocrit. These sources of variation have impeded an accurate determination of a sample’s equivalent input plasma volume, which is necessary to calculate plasma concentration from a measurement of the amount of analyte in the DBS punch. In response to this perceived need and the recent uptick in interest in wider application of dried blood samples, a number of collection devices are being developed that aim to reduce this blood volume uncertainty through volumetric measurement prior to sample drying, and ultimately perhaps achieve plasma volume precision as well through use of an integrated plasma separation step. We have addressed this issue in a different way via a personalized normalization strategy that is compatible with both existing DBS cards and novel devices. To do this we employ a normalization method based on measurements of a small set of plasma proteins that remain relatively unchanged over time in an individual, and show that this strategy significantly reduces the variation observed in other proteins in sets of DBS samples. The result is a system that provides blood protein measurements with precision comparable to conventional clinical assays performed on standard venipuncture samples.

To measure panels of selected blood proteins in DBS we used stable isotope standards and capture by anti-peptide antibodies (SISCAPA; [5,8]). This technology involves tryptic digestion of samples followed by immuno-affinity enrichment of surrogate proteotypic peptides coupled with peptide quantitation using liquid chromatography-multiple reaction monitoring (LC-MRM) mass spectrometry [9]. Tryptic digestion overcomes the effects of sample drying and essentially converts large molecules (proteins) into smaller peptides amenable to precise MRM quantitation, while SISCAPA peptide enrichment eliminates most sample matrix, improves sensitivity and throughput and enables measurement of proteins present in widely disparate (>10^6^-fold) concentrations in the same multiplex assay. In the work reported here we measured a series of 26 biomarker proteins selected to track multiple aspects of human health and disease, including inflammation, coagulation, lipoprotein metabolism, kidney function and iron metabolism. In addition, several selected proteins were measured as surrogates for blood cell counts (erythrocytes, leukocytes and neutrophils, the most abundant class of leukocytes]. Our targeted approach focuses on precise measurement of selected proteins with known clinical significance [4] rather than a broad survey covering hundreds of candidates [10–13] (the objective of classical biomarker proteomics), thereby improving throughput, precision, robustness, sensitivity and cost at the expense of potential for *de novo* biomarker discovery. The precision and throughput of this approach facilitate detection of small quantitative changes in large sample sets, thereby enabling high-frequency longitudinal biomarker studies as described here, while the robustness and sensitivity of established clinical applications [14,15] provide a straightforward path to clinical implementation.

Inflammation events provide perhaps the best illustration of a broad pattern of recurring large-amplitude biomarker variation in humans. In clinical contexts, inflammation is usually measured using the blood biomarker C-reactive protein (CRP; [16]) a key component of the adaptive immune system. Quantitative tests for CRP are widely used in laboratory medicine, and the results have clinical significance over a large dynamic range: increases of more than 100-fold occur in major infections [17], while much smaller increases (<2-fold) are associated with elevated cardiovascular disease risk [18]. Use of any single biomarker for such a complex system relies on the assumption that inflammation is a relatively homogeneous response governed by familiar regulatory mechanisms [19] and operating similarly in most individuals in response to inflammatory triggers. In fact, the concentrations of many other plasma proteins also change in response to inflammation, including cytokines (short-lived, low-abundance proteins regulating inflammatory responses) and a broad set of acute phase response (APR; [20]) proteins associated with many functions. While cytokines such as interleukin-6 (IL-6) are occasionally used as clinical diagnostic tests and have been measured in DBS [21], they typically appear briefly, serving as signals initiating an inflammatory process [22], and at very low concentrations (low pg/mL; ∼ 10^5^-fold lower abundance than APR proteins). In this manuscript, we focused on the APR proteins to track inflammatory events with timescales ranging from days to months. These included serum amyloid A (SAA; see Table 2 for complete list of protein abbreviations), LPS-binding protein (LPSBP), mannose binding lectin (MBL), alpha-1-acid glycoprotein (A1AG; orosomucoid), haptoglobin (Hp), fibrinogen (FibG) and complement C3 (all of which are components of the positive APR whose plasma levels increase in response to inflammation) and albumin (Alb), transferrin (Tf), hemopexin (Hx) apo A-I lipoprotein (ApoA1), and IgM (which are components of the negative APR whose plasma levels decrease in response to inflammation).

Increases in CRP are observed in connection with infections [23], arthritis [24], surgery [25], sleep apnea [26], depression [27], Parkinson’s disease [28], pregnancy [29], inflammatory bowel disease [30] and cancer [31], to name a few examples. In each of these cases, increased inflammation is associated with poorer outcome, but the increase (as currently measured via CRP) is not disease-specific. Because of this lack of disease specificity and the assumption that CRP alone is sufficient to track inflammation, few data are available on the detailed dynamics of multiple APR proteins in specific disease situations or in individuals outside small research studies. To obtain a more complete picture, we selected the panel of APR proteins that includes molecules involved in both innate (SAA, CRP, LPSBP, MBL) and adaptive immune responses (IgM) and proteins involved in drug transport (Alb, A1AG), iron scavenging and transport (Hp, Hx), the complement cascade (C3), coagulation (FibG) and lipid metabolism (ApoA1) to monitor inflammation’s impact on multiple systems, of interest on timescales from acute to chronic.

The results presented here illustrate the complex and informative behavior of APR proteins in tracking inflammation due to multiple causes over many orders of magnitude. Most important, the data reveal the practicality of major improvements in diagnostic test interpretation using personalized baseline values and the feasibility of personalized multiparameter biomarker response models useful in tracking a variety of planned health interventions, e.g., clinical trials.

## Results

### Human Blood Samples, SISCAPA analysis and DBS sample normalization

A unique set of 1,522 longitudinal DBS samples was collected on Whatman 903 cards by 8 participants between 2008 and 2017 and included extensive periods of daily sampling (Table 1). Biomarker proteins selected for clinical relevance to inflammation and chronic disease (Table 2; including protein abbreviations used throughout) were measured in punches taken from the DBS cards using an automated, multiplexed SISCAPA protocol [5,32] in three tranches as described in Materials & Methods. Protein amounts were determined by mass spectrometry and expressed as femtomole per sample punch.

**Table 1.** Description of samples

| Subject | Age (yr) | Sex | Overall health | 2008 | 2009 | 2010 | 2011 | 2012 | 2013 | 2014 | 2015 | 2016 | 2017 | Total samples | Span (days) |
| --- | --- | --- | --- | --- | --- | --- | --- | --- | --- | --- | --- | --- | --- | --- | --- |
| S-01 | 62 | F | Good |  | 7 | 17 |  |  |  | 24 | 17 | 54 | 67 | 186 | 2,791 |
| S-04 | 71 | F | Good |  |  |  |  |  | 10 | 15 | 43 | 52 | 130 | 250 | 1,647 |
| S-07 | 55 | M | Good |  |  |  |  |  | 18 | 31 |  |  |  | 49 | 393 |
| S-10 | 37 | M | Crohn's |  |  |  |  |  |  | 49 | 52 | 183 |  | 284 | 852 |
| S-17 | 32 | M | Good |  |  |  |  | 2 | 28 | 19 | 19 | 19 |  | 87 | 1,415 |
| S-18 | 69 | M | Good | 14 | 12 | 6 | 52 | 22 | 31 | 50 | 52 | 48 | 124 | 411 | 3,292 |
| S-20 | 47 | F | Good |  |  |  |  |  | 21 | 24 |  |  |  | 45 | 390 |
| S-22 | 72 | M | Good |  |  |  |  |  | 99 | 29 | 40 | 42 |  | 210 | 1,224 |
| Total |  |  |  | 14 | 19 | 23 | 52 | 24 | 207 | 241 | 223 | 398 | 321 | 1,522 | 12,004 |

**Table 2.**
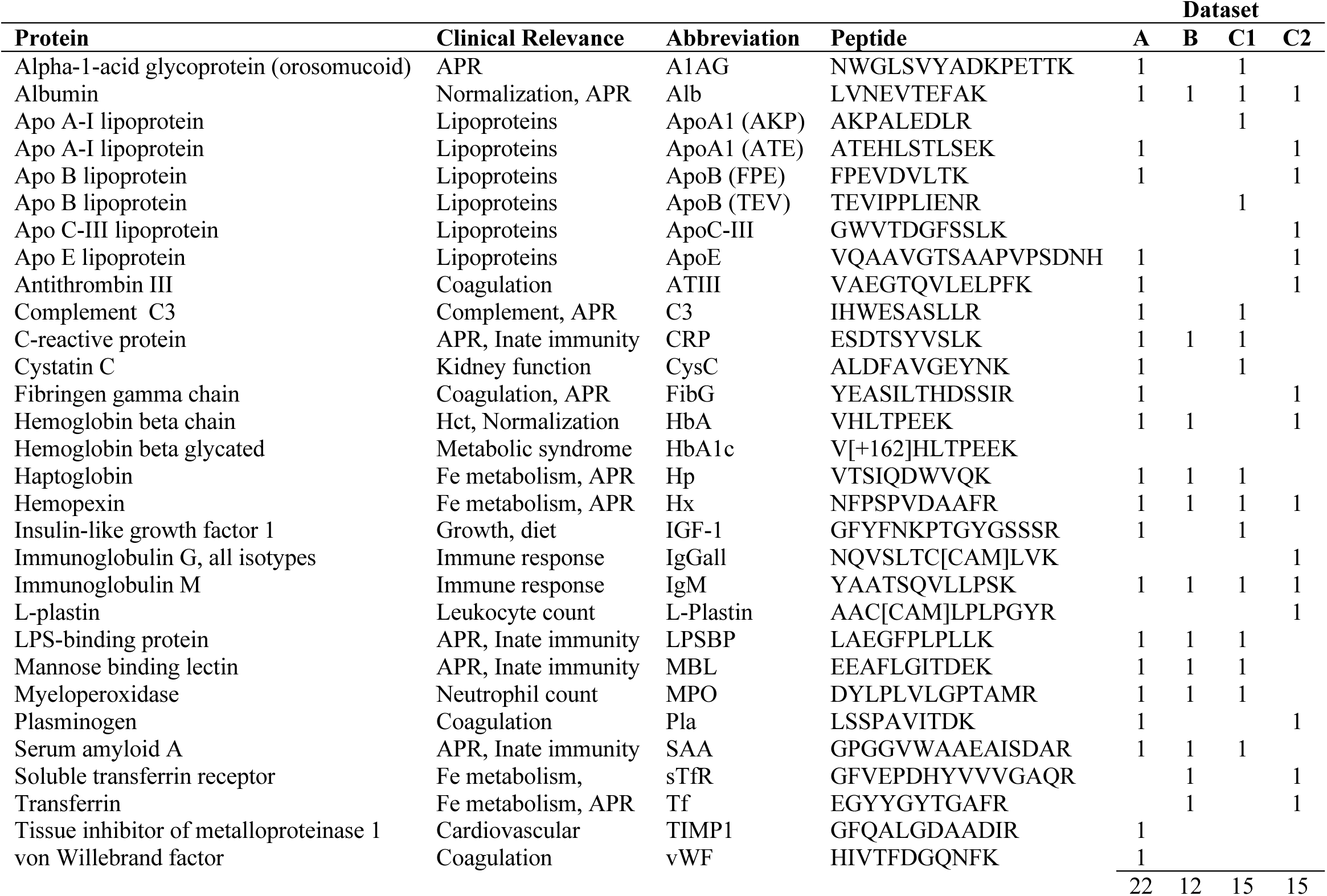
Proteins and peptides measured in the 3 datasets. A single proteotypic tryptic peptide (sequence unique to the target) was measured as a surrogate for each protein, except for ApoA-I and ApoB lipoproteins for which 2 peptides were measured in some cases. Columns A, B and C refer to the three datasets (C1 and C2 are the two sequentially-measured multiplex panels into which the Dataset C panel was divided), and a value of 1 indicates the protein was measured in the corresponding dataset.

Variations in the volume of blood sampled in a DBS punch have historically limited the precision of DBS measurements of biomarkers. Theoretically this volume is ∼14 μL of whole blood in the case of a 1/4” (∼6mm) diameter circular punch from a Whatman 903 card, but in practice this amount can vary by +/-10-15% due to a number of factors such as hematocrit, viscosity and chromatographic effects that are often poorly controlled [33]. In order to overcome this limitation, we developed a novel method for normalizing the blood volume in longitudinal DBS punches from an individual, with the objective of maximizing assay precision when comparing serial samples against personal baselines. The method uses a sample-specific scale factor computed from an equally-weighted combination of three proteins (albumin, hemopexin and IgM) whose abundances are usually very stable within individuals over time and which, when they do vary under extreme circumstances (e.g., major inflammatory events), typically counterbalance one another. The calculated scale factors applied to individual longitudinal DBS spots in this study fell in the range of 0.7-1.3 as expected, with an overall CV of 16.2%. Duplicate measurements of the scale factor in Dataset C (determined when measuring the first plex *vs* the second plex - i.e., tryptic digestion of a single sample followed by separate sequential SISCAPA captures and LC-MRM measurements) showed very good agreement and were related by a slope of 0.99 and R^2^ of 0.98.

### Precision of SISCAPA-LC-MRM measurements in dried blood spots

Replicate standard samples dried on Whatman 903 filter paper were included in each 96 well plate processed for SISCAPA measurement, normalized using the above methods and used to evaluate the precision of each assay in each dataset. The average coefficient of variation (CV) across the assays used in Datasets A, B and C were 7.4%, 6.0% and 4.9% respectively, consistent with expectations for clinically useful assays. CVs were typically somewhat higher in Datasets A and B, which were analyzed earlier in the process of assay optimization. In Dataset C the CV over all 26 peptides measured in two successive multiplex enrichments from 4 replicate dried blood standards in each of nine 96-well plates (72 measurements) was reduced from 10.2% prior to volume normalization to 4.6% after normalization (Supplementary Table 1). Similarly, the overall average CV in Dataset A was reduced from 9.4% to 7.4% by normalization. In Dataset B, normalization had little effect (6.5% vs 6.1%) because in this case the standards were prepared by drying the same measured volume of whole blood onto pre-cut Whatman 903 filter paper disks in wells, such that there was little or no volume variation to correct. In Dataset C, a number of proteins including apolipoprotein A1 (Apo A-I) and apolipoprotein B (Apo B), insulin-like growth factor 1 (IGF-1) and transferrin, in addition to the normalizing triad (albumin, hemopexin and IgM), yielded CVs of ≤ 2.5% in the replicate standard samples. Key inflammation markers such as C-reactive protein (CRP), serum amyloid A (SAA) and lipopolysaccharide binding protein (LPSBP) showed normalized CVs in the standard samples of 3.3% −3.7%. The positive effect of volume normalization significantly smoothed plots of most proteins across longitudinal sample sets as expected (Supplementary Figure 1).

The sample sets contributed by the 8 subjects each produced a tight cluster of points in a plot of IgM vs. Hx (Figure 1), demonstrating the stability of both the DBS samples and the analytical approach. The 8 subjects can be clearly differentiated from one another using only these two parameters and in fact most of the APR proteins measured showed significant inter-individual difference (Supplementary Figure 2). Subjects S-04 and S-18 (of opposite sex, contributing 411 and 250 samples respectively) share a living environment but their biomarkers appear as completely separated clusters, indicating that most of this between-subject variation observed is not determined by environment at the time of collection. In general, points diverging from the cluster centroids are associated with major inflammation events.

**Fig. 1.**
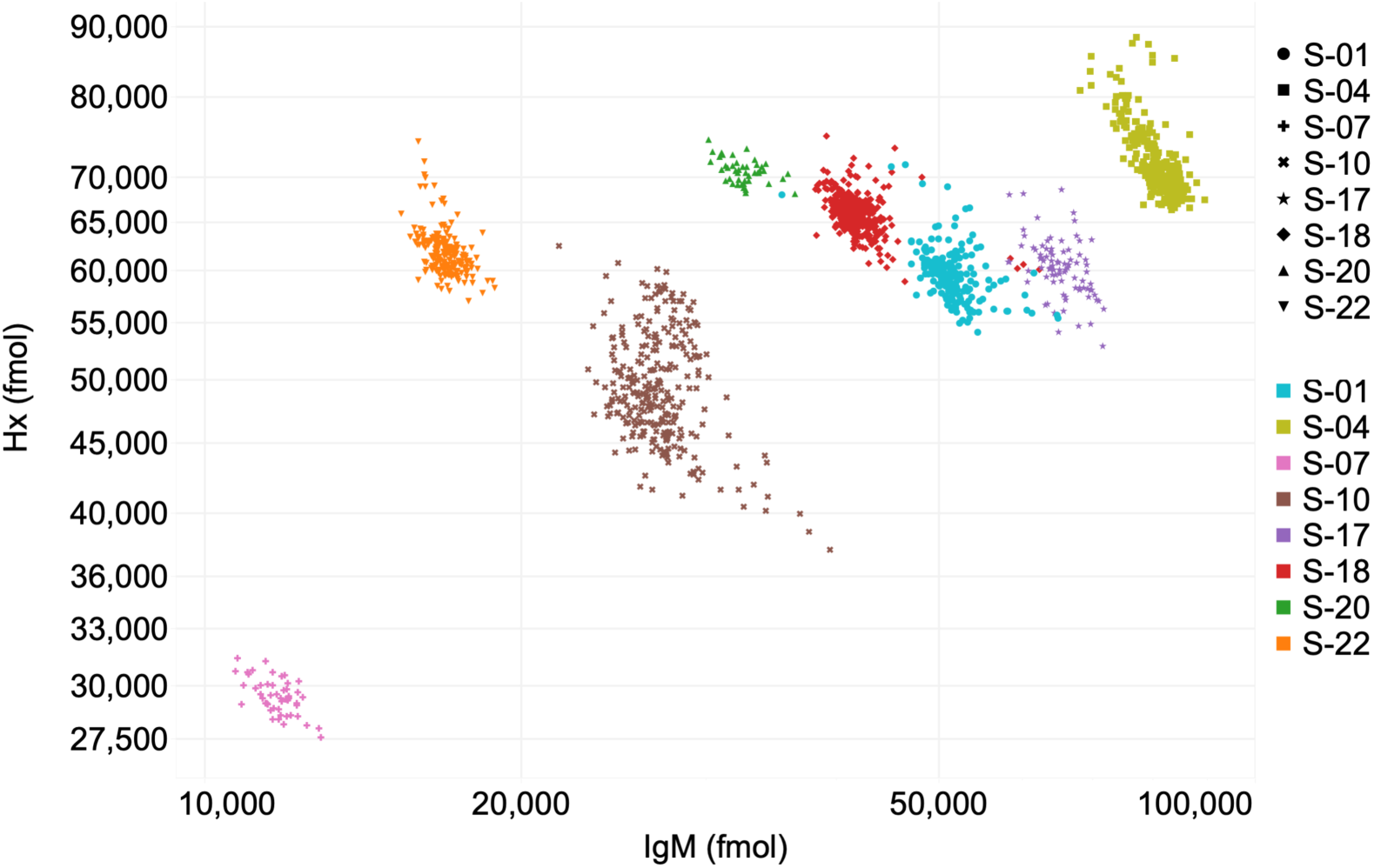
A plot (log-log) of IgM vs Hx values for all 1522 samples, marked by subject.

### Personalized baselines and standard deviations

The largest perturbations observed in most proteins occurred during inflammatory events that were noted by the study subjects and were coincident with increases in the acute phase reactants CRP and SAA. In order to calculate personal “normal” baseline levels for each biomarker protein that excluded inflammation events, we identified, for each subject, the subset of samples with CRP below a cutoff that was set at the personal median CRP value across all of a subject’s samples; i.e., selecting the half of each subject’s samples with lowest CRP values. For each protein, the average value in these low-inflammation baseline samples was taken as the subject’s personal baseline and the standard deviation in these samples allowed calculation of a personal baseline CV. The average baseline CV over all subjects, samples and proteins was 12.3% (12.4%, 17.0% and 10.2% in Datasets A, B, and C respectively) compared to 38.0% (39.2%, 68.7% and 31.3% in Datasets A, B, and C) for combined baseline and non-baseline (i.e., all) samples (Table 3). On average, the ratio of assay CV (from replicate standards) to average subject baseline CV (from subjects’ longitudinal samples) was 0.52, satisfying the criterion (<0.6) allowing statistically meaningful analysis of within-subject biological variation [34]. In subject S-22 CRP and SAA showed maximum increases of up to 497- and 1,062-fold (equivalent to 2,673 and 3,688 personal standard deviations) respectively, above personal baselines, illustrating the >1,000-fold dynamic range of statistically significant (>2 SD) within-person changes.

**Table 3.**
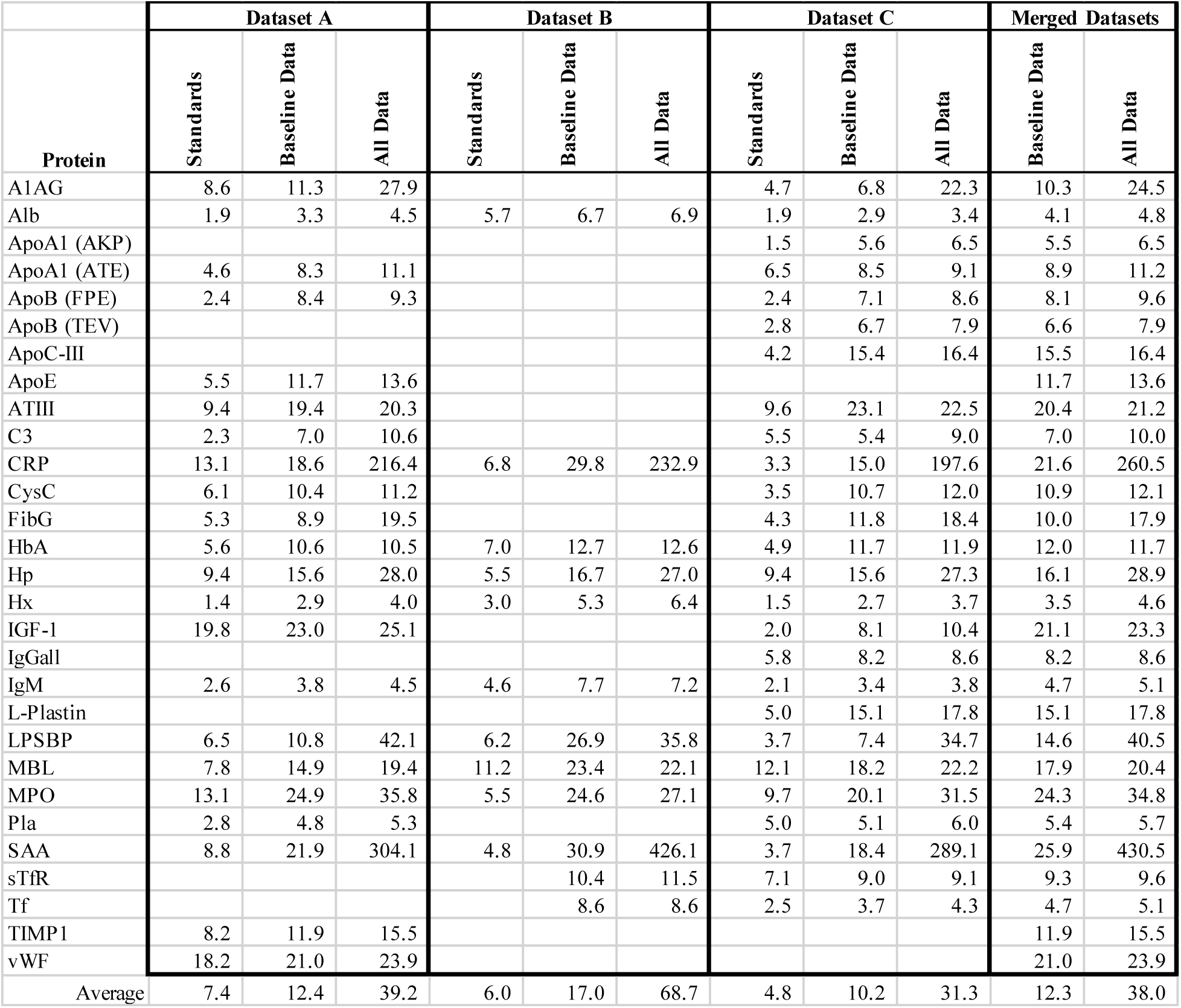
Coefficients of variation of protein measurements (%CV) in all three datasets separately and in combination (merged). CVs in Standard samples (workflow analytical CV), subject baseline data (averaged over subjects) and all subject data (including inflammation events) are separately tabulated.

### Correlations among inflammation biomarkers

Protein:protein correlation matrices were calculated to help uncover relationships among the inflammation–associated biomarkers. Figure 2 shows the correlation matrix obtained using all 1,522 subject samples after volume normalization (using Alb, Hx, IgM) and an additional normalization step (division by each subject’s personal median value for each protein) to reduce the impact of inter-individual differences. A set of proteins at the upper left (Area A: CRP, LPSBP, SAA, A1AG, Hp, C3, MBL and FibG), together with normalizing protein Hx on the right, exhibit strong positive mutual correlations: all of these are classical positive acute phase reactants. A set of strong (Alb, Tf, ApoA1 and ApoC-III) and weaker (IgM and IgGall) negative acute phase reactants are anti-correlated with the positive APR (particularly with Hp, whose slow time course most closely resembles the slow time course of the negative APR). The observed correlations among acute phase proteins are common to the individual subjects (Supplementary figure 3) and feature very close correlations between CRP (the primary clinical inflammation marker) and SAA (pairwise correlation of 0.89 across all the data) or LPSBP (pairwise correlation of 0.84).

**Fig 2.**
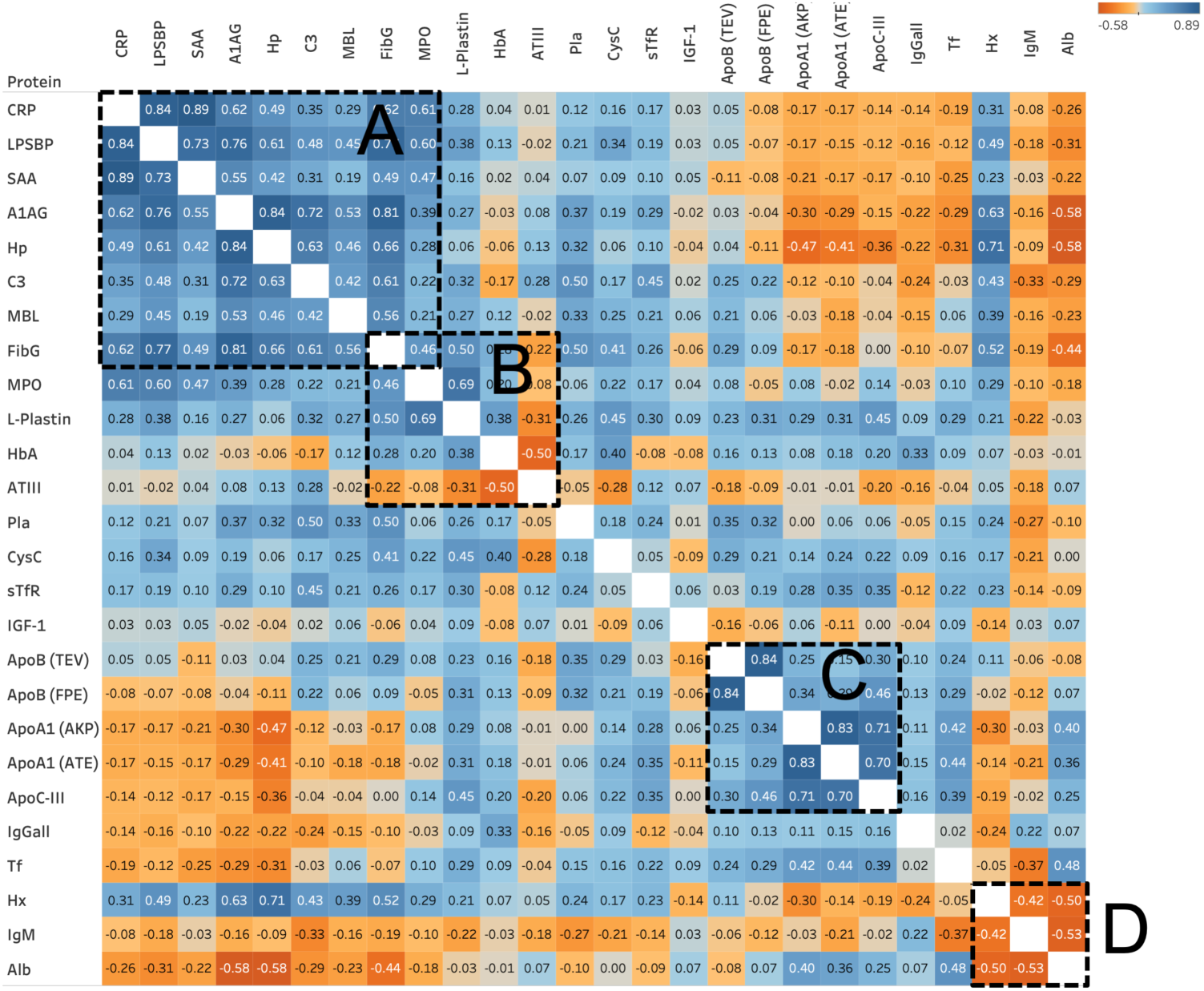
Protein:Protein correlation matrix. A matrix describing correlation of variation in each protein to the others, calculated over all samples from all subjects after sample volume normalization and division by personal baseline average values (to minimize the impact of subject:subject differences in protein amount). A): acute phase response proteins; B): cluster related to coagulation and blood viscosity; C) lipoproteins; D) proteins used in volume normalization.

Similar correlation results were obtained using only the half of each subject’s samples with the lowest CRP (i.e., in the baseline low-inflammation data; Supplementary figure 4), except that correlations and anti-correlations among the acute phase reactants are significantly reduced due to the removal of all prominent inflammation events from the data. Nevertheless, the correlations between CRP and SAA or LPSBP remained positive and significant in the baseline (low CRP) samples (0.30 and 0.31 respectively), clearly demonstrating that low-level fluctuations in these proteins contain biological signal. This persistent correlation (indicative of coregulation) among the acute phase proteins in the low-inflammation data appears clearly in scatterplots plots of CRP vs SAA and LPSBP (Figure 3, showing data from the five subjects who contributed most samples). During inflammatory events, where CRP varies by 100-fold, the SAA response was consistent across subjects, increasing by up to 1,000-fold over baseline. LPSBP levels increased similarly, but the observed increase was less than 10-fold at maximum.

**Fig 3.**
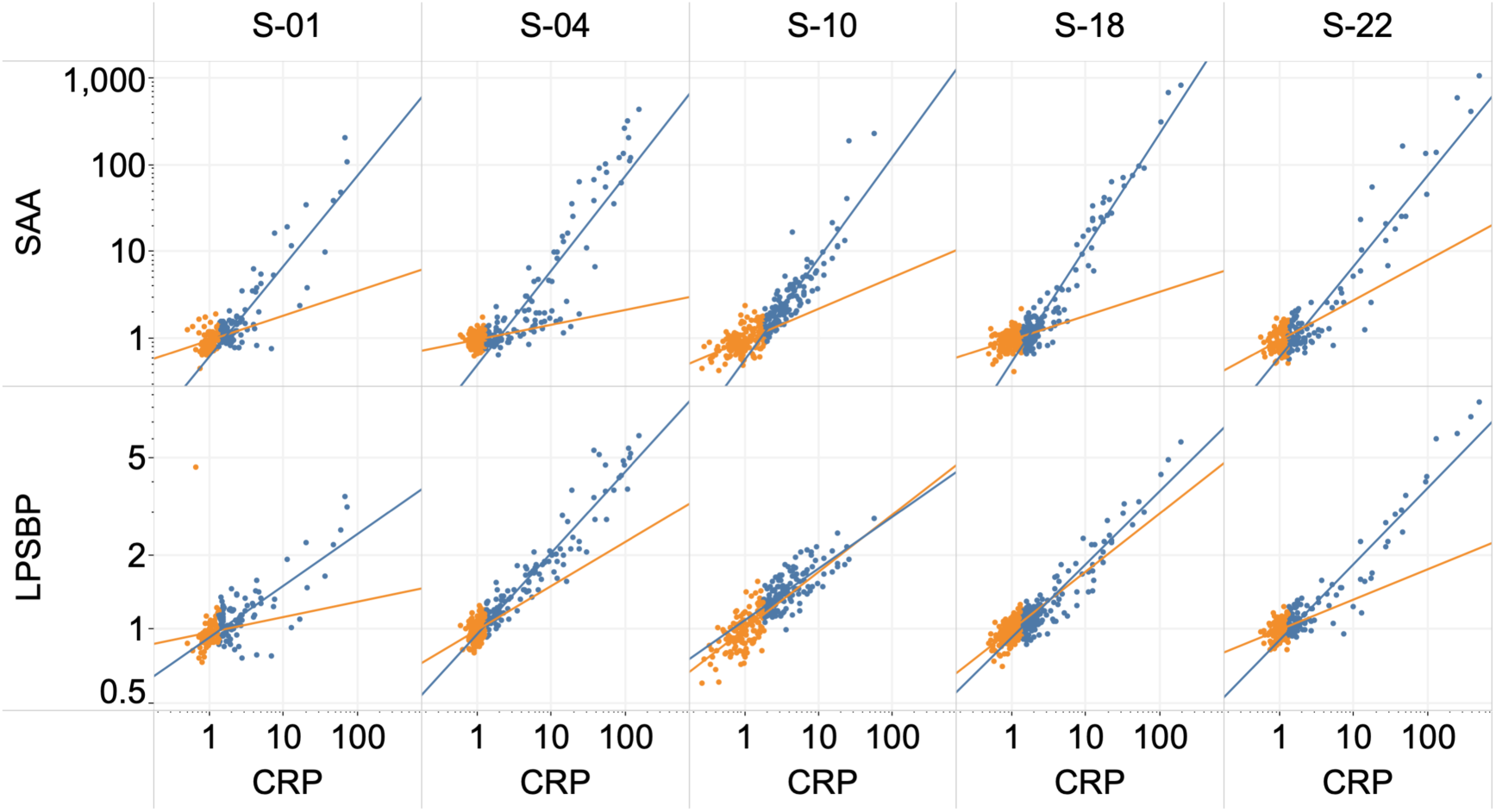
Relationship between CRP and SAA or LPSBP. Scatterplots (log-log) relating CRP to SAA or LPSBP in the individual samples (dots) contributed by 5 subjects. Baseline samples (the half with lowest CRP) are shown in orange, and the higher half of samples in blue. All values were normalized by the personal average of the baseline samples.

Other large-scale features of the correlation data (Figure 2) include a second block of correlations (B) comprised of AT-III, HbA, L-plastin, MPO and FibG, in which AT-III is negatively correlated with the other members. A third block (C) is comprised of the apolipoproteins (Apo A-I, Apo B and Apo C-III), in which the measurements of pairs of peptides from the same protein (i.e., Apo B TEV and FPE peptides, or Apo A-I AKP or ATE peptides) showed the expected very strong correlations (0.79 and 0.76 across all subjects and samples). A fourth block (D) is comprised of the triad Alb, IgM and Hx, whose mutual anti-correlation is primarily driven by their use in normalizing plasma volume in the DBS samples.

### Inflammation events

Among the 5 subjects for which extended (>150) longitudinal sample series were available, we identified 58 inflammation events in which CRP, the primary clinical indicator of inflammation, was increased significantly above baseline levels in multiple samples collected over a brief interval allowing identification of a temporal maximum (Figure 4). Of these events, about half (26 events) included at least one sample in which CRP was increased by more than 10-fold above baseline and all were confirmed by coordinated increases in CRP, SAA and LPSBP. In contrast to the large increases seen in SAA and CRP in these events, the remaining APR proteins (LPSBP, A1AG, Hp, MBL, FibG, C3 and Hx) showed much smaller effects, with fold-increases relative to baseline ranging from 0.036 (LPSBP) to 0.002 (Hx) as great as the fold-increase in CRP. Inflammation events occupied a significant proportion of the total sample series: the proportion of samples in which CRP is more than 2-fold higher than the personal baseline ranged from 12-45% among these 5 subjects (Supplementary Table 2).

**Fig 4.**
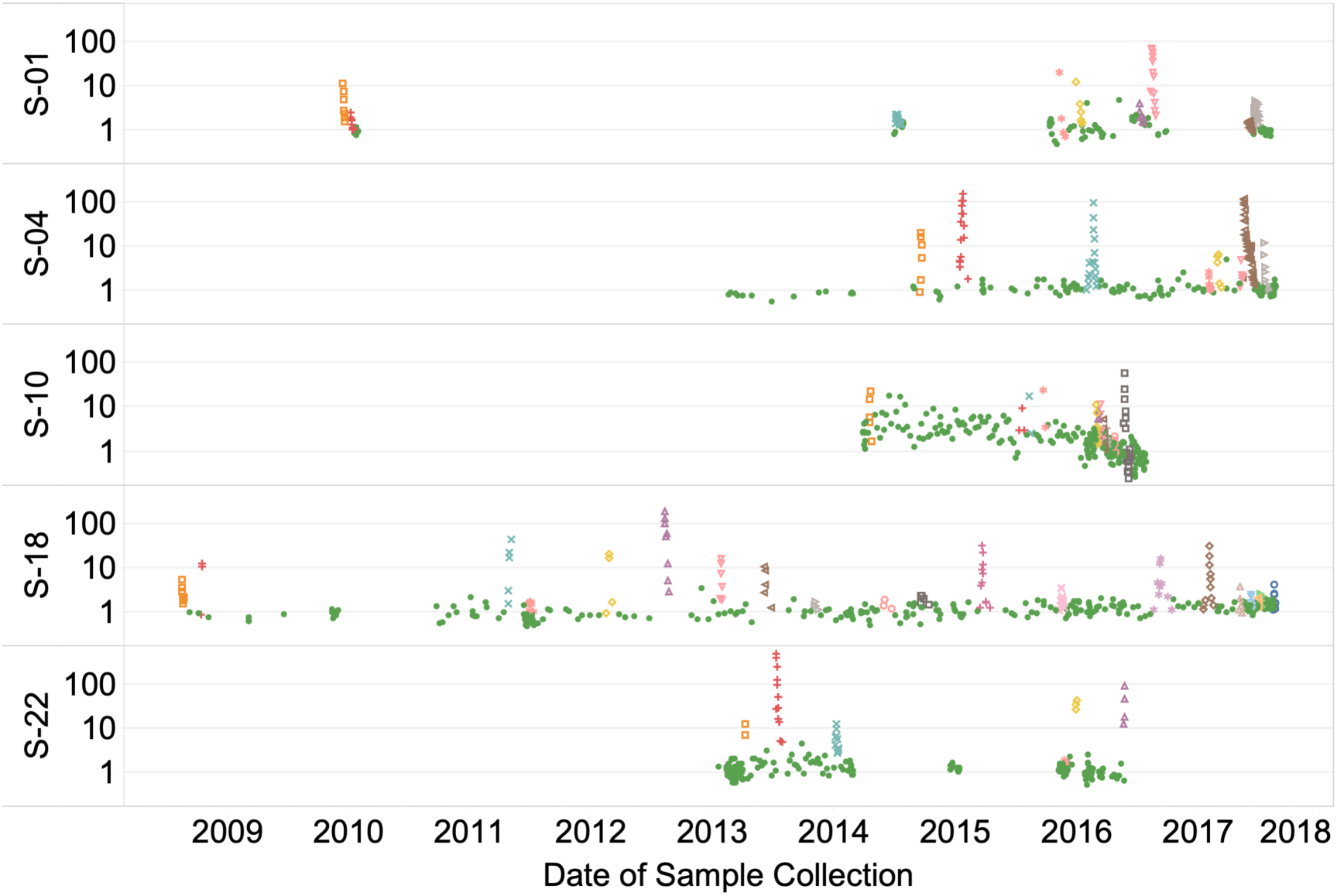
CRP inflammation events. Inflammation events identified among the CRP measurements (y-axis; log scale) for 5 subjects. Event samples are distinguished by different colors and symbols; remaining samples are shown as green dots.

Among the events captured in these samples there are clear examples of at least three causes of acute inflammation.

### Surgery

Subject S-04 underwent an elective total hip arthroplasty involving a short (∼2 hr) surgical intervention and subsequent recovery period of 97 days during which daily (or more frequent) samples were collected (Figure 5). Following an initial rise immediately following surgery, inflammation markers (here plotted on log scales), declined at different rates until day 55, by which point the levels approached a new baseline approximately 50% below the pre-surgery level. The pre-surgery levels were somewhat higher than those in the immediately preceding period, possibly due to the required suspension of analgesics before surgery. CRP and SAA were induced much more strongly than other inflammation indicators (with maximum inductions of 113- and 136-fold respectively). The magnitudes of the observed responses (in fold-change from personal baselines) were: SAA > CRP >> LPSBP >FibG, Hp, A1AG > MBL, C3 > Hx (136, 113, 5.4, 3.0, 2.9, 2.7, 2.0, 1.8, and 1.2-fold respectively), overall a ∼500-fold range of responses relative to personal baselines. In the period between days 8 and 30, a series of post-surgical jumps in SAA and CRP (e.g., days 9, 18 and 22) indicated renewed inflammatory activity that generally coincided with increased requirement for pain medication and subject reports of strain in the surgical area, followed by smooth declines to baseline.

**Fig 5.**
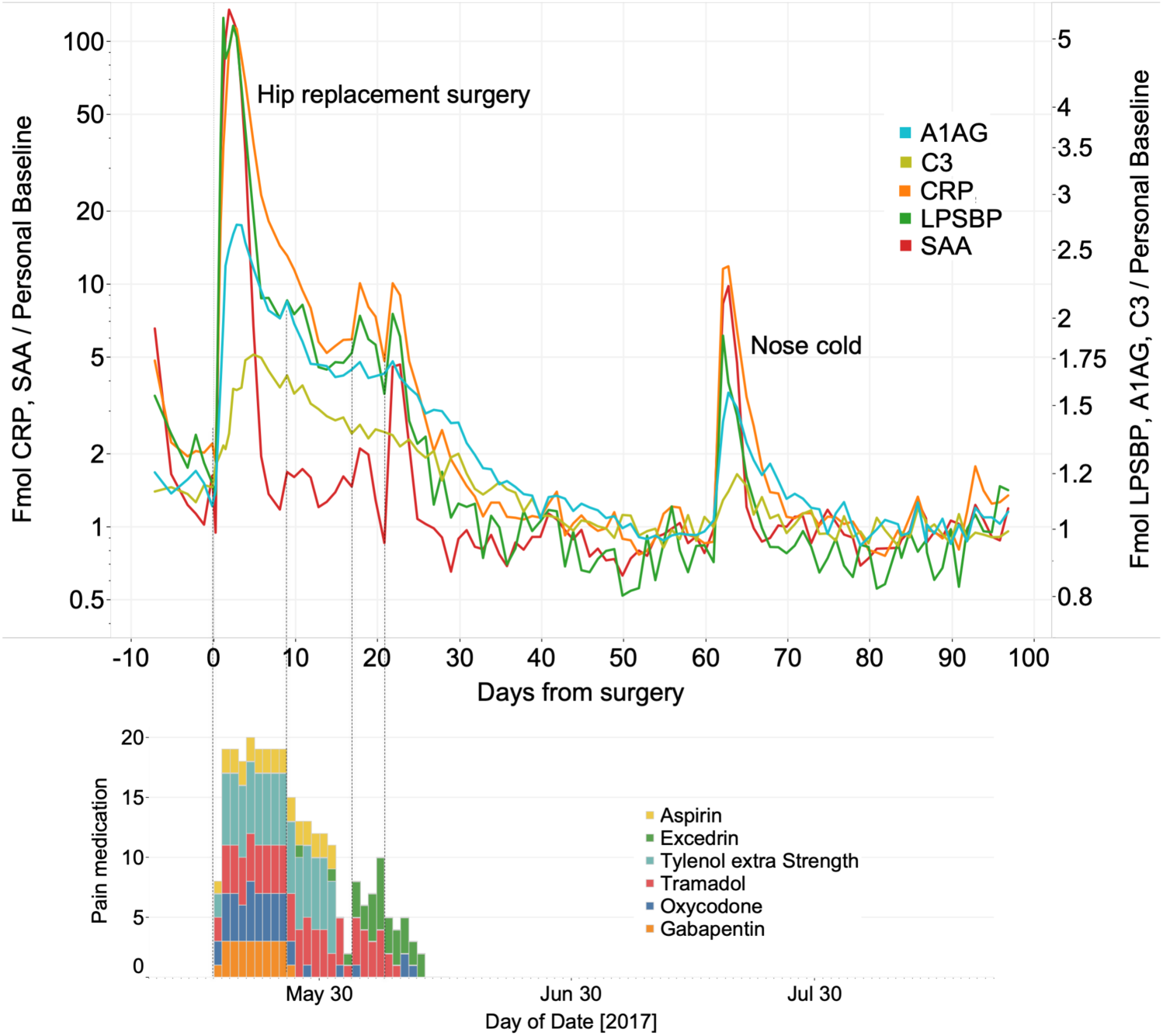
Inflammation related to surgery. Amounts of 5 inflammation proteins (log scales) over 100 days including a total hip artheroplasty and an upper respiratory infection (upper panel) and requirement for pain medication associated with the surgery (lower panel).

Figure 6 shows the time course after surgery of 9 acute phase proteins normalized to the same peak response and smoothed in order to reveal relative peak times. LPSBP achieved its peak level first (∼1.2 days post-surgery), followed by SAA, CRP, A1AG and MBL, Hp, Fib G, Hx and C3, with the last peaking approximately 5 days post-surgery.

**Fig 6.**
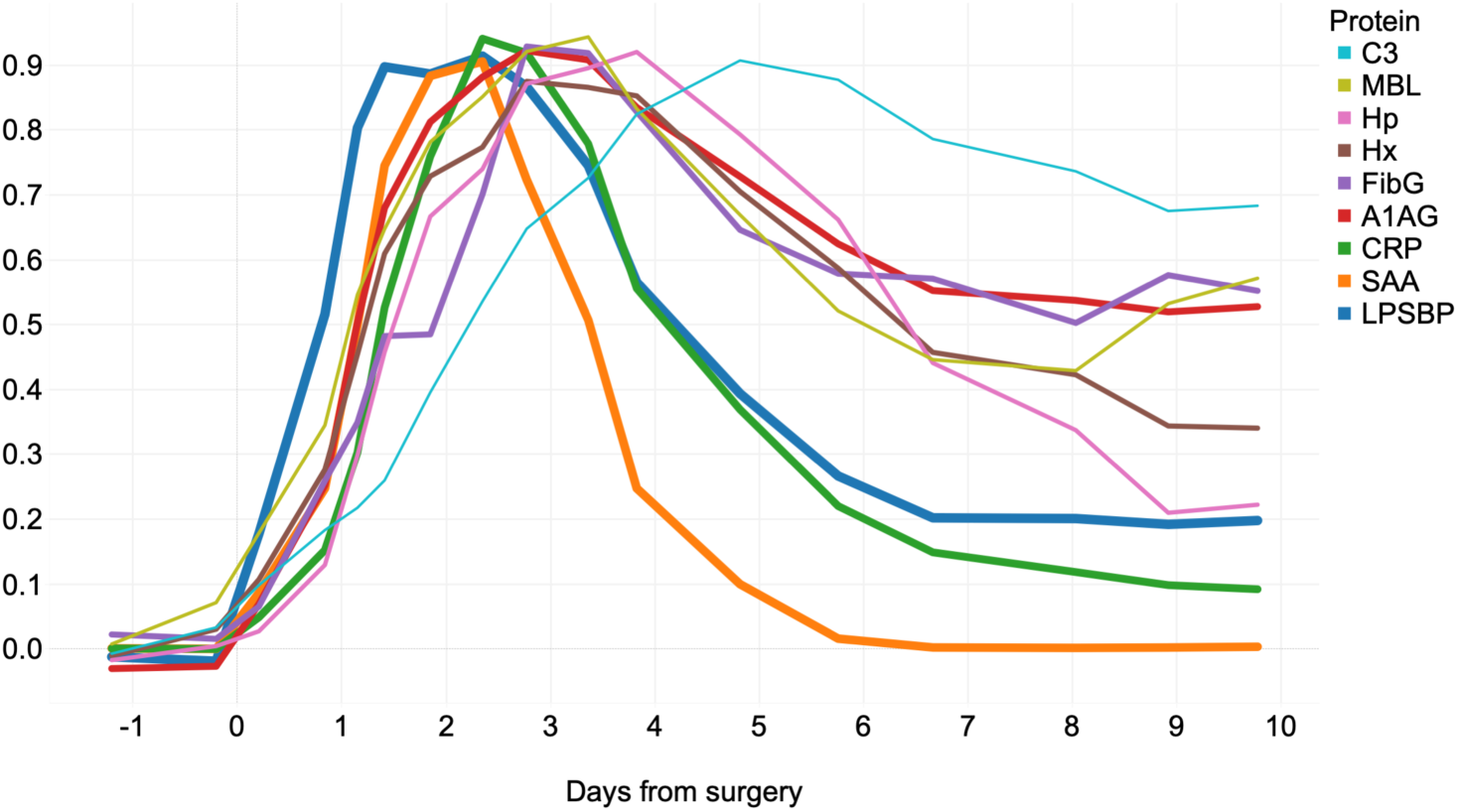
Relative timescales of inflammatory protein responses to surgery. Amounts of 9 inflammation proteins over 10 days including a total hip artheroplasty, with each protein normalized to approximately the same scale of change in order to illustrate differences in timecourse.

The difference in timing between earlier and later-peaking inflammation markers can be used to visualize an inflammatory response trajectory in two dimensions, from baseline, through injury and returning to baseline [35]. Figure 7 shows such a trajectory plotting SAA (fast response, up to 137-fold above baseline) vs. Hp (slow response, maximum at 1.77-fold above baseline) in 85 successive DBS samples over 80 days. The large red loop tracks progress (counterclockwise) from surgery (indicated by a yellow +) through initial healing (day 8), followed by a period of re-inflammation (days 10-27; orange, coincident with increased use of pain medication) and recovery to baseline (day 60; blue).

**Fig 7.**
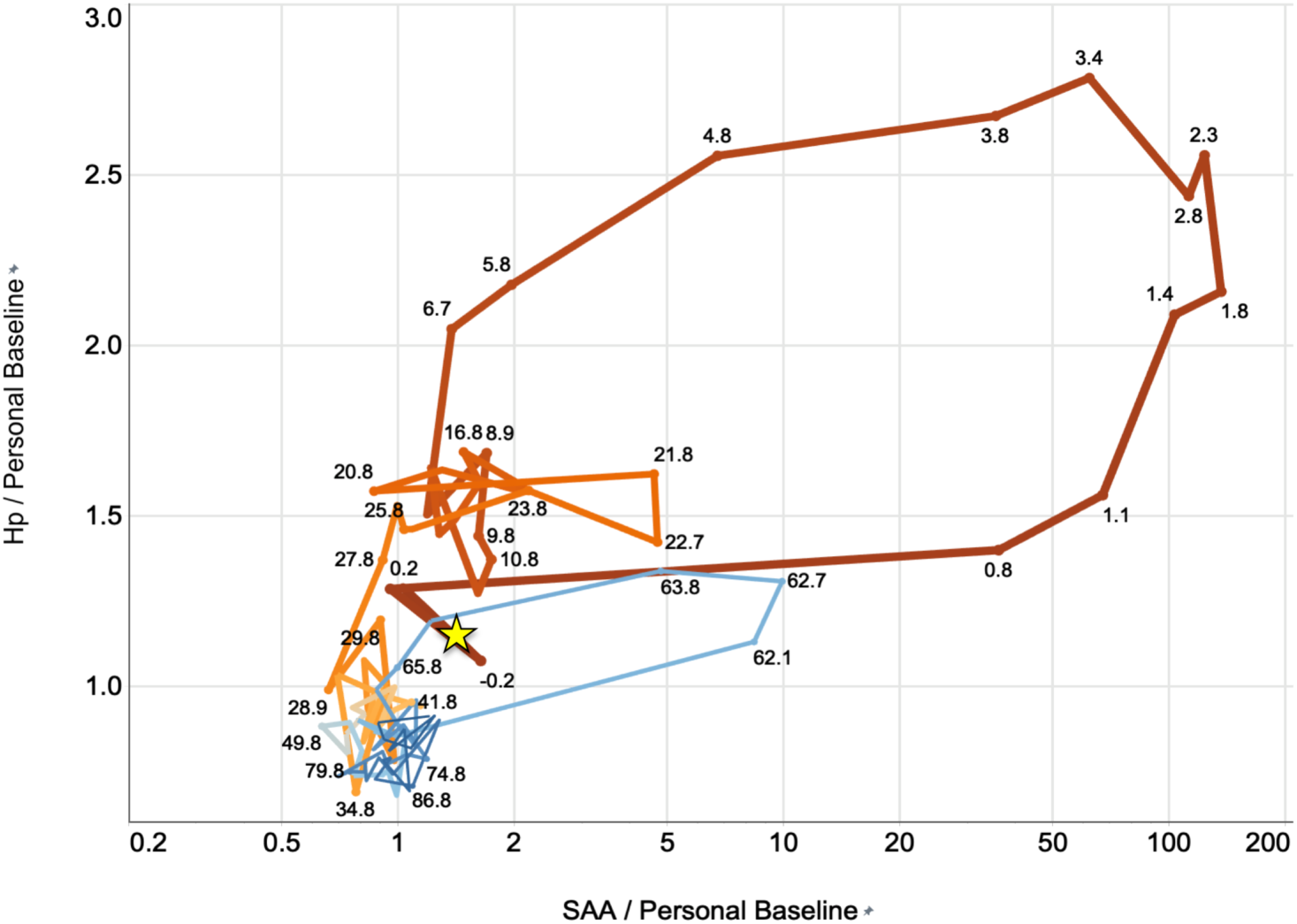
Loop plot of inflammation timecourse. A plot of SAA vs Hp (each normalized by their respective average values in baseline samples, SAA on log scale, Hp linear), with successive time points linked by lines to form loops. Time is represented by line color (deep red at the time of surgery grading smoothly to blue 75 days later), a steady decrease in line width over time, and by numbers (days post-surgery). The time of surgery is indicated by a yellow star.

### Infection

Infections of various kinds constituted a second major driver of inflammation events. Figure 5 includes such an event (an upper respiratory tract infection perceived by the subject as a “nose cold”) on day 62 following surgery. While the overall magnitude of the biomarker response in this case was roughly 10% as great as the response to surgery, the kinetics of the relative increases of the 5 proteins shown were similar, rising rapidly within 1-2 days and declining to baseline within 10 days. In Figure 7, the infection appears as a response trajectory in the bottom center of the plot (blue loop beginning and ending in the cloud of points at the baseline; also traversed counterclockwise as described above).

Figure 8 shows loop and time course plots for 7 significant infections occurring in 5 subjects, each involving SAA increases >100-fold from personal baseline median values. These events were reported by subjects as: respiratory infection (S-01 E7; i.e., event 7 in subject S-01); respiratory infections (S-04 E2, E3 and E7); food poisoning (S-10 E11); pneumonia (confirmed by X-ray) (S-18 E6); and kidney infection (S-22 E2). None of the infections required hospitalization, though in some cases antibiotics were administered. The events show a striking consistency among subjects in the maximum relative induction levels of SAA in major infections, but significant differences in the time evolution of the responses (i.e., differences in the shape of the response loops). Subject S-18, with the largest sample series, exhibited a number of similar smaller loops (below the large event 6 loop) that were described as “nose colds” and have a remarkably consistent size and structure, likely indicating a reproducible response to similar infectious agents (e.g., rhinoviruses).

**Fig. 8.**
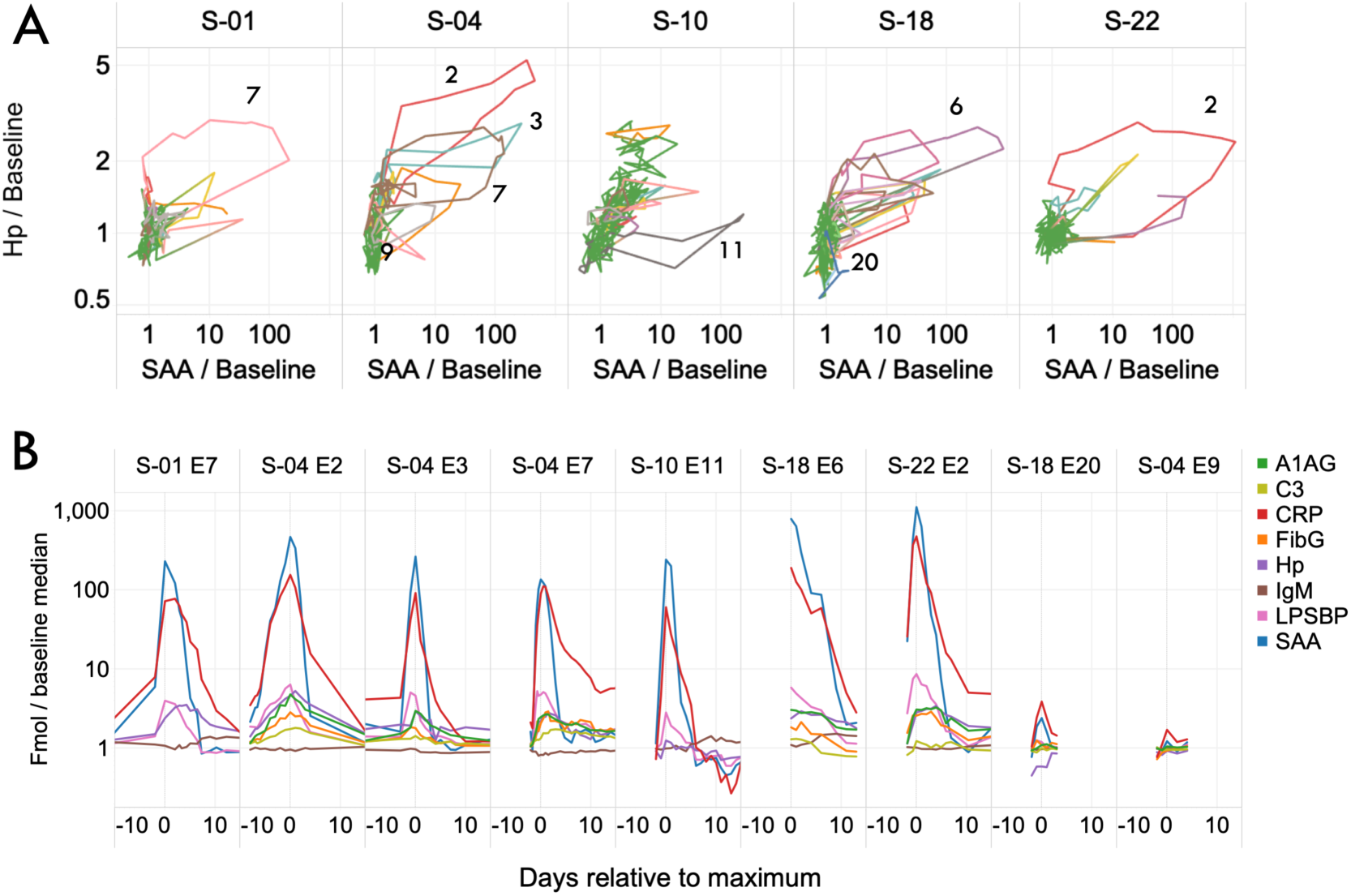
Loop plots and timecourses of major inflammation events. Seven major inflammation events and two influenza vaccinations in 5 subjects involving SAA increases >100-fold from baseline. A) SAA vs Hp counterclock-wise loop plots (log-log). Samples not consider part of an event are plotted in green. Numbered loops in various colors correspond to events in B, which show the timecourses of 8 acute phase proteins (in fold-change from personal baseline average, log scale) as a function of days from maximum SAA.

Despite the high correlation between CRP and SAA responses, the relationship between the maximum levels achieved by CRP and SAA levels was clearly non-linear. An examination of the set of events (Figure 9A) in which a local maximum was determined (i.e., for which samples were available with lower levels just before and after a maximum) shows that SAA’s maximum induction relative to baseline exceeds CRP’s for the highest level events, but CRP achieves higher levels of induction than SAA for the smaller events. This relationship can be modeled by a power fit (Figure 9B) in which SAA is related to CRP raised to a power between 1.31 and 1.52, except for subject S-10 for which the power is 1.75.

**Fig 9.**
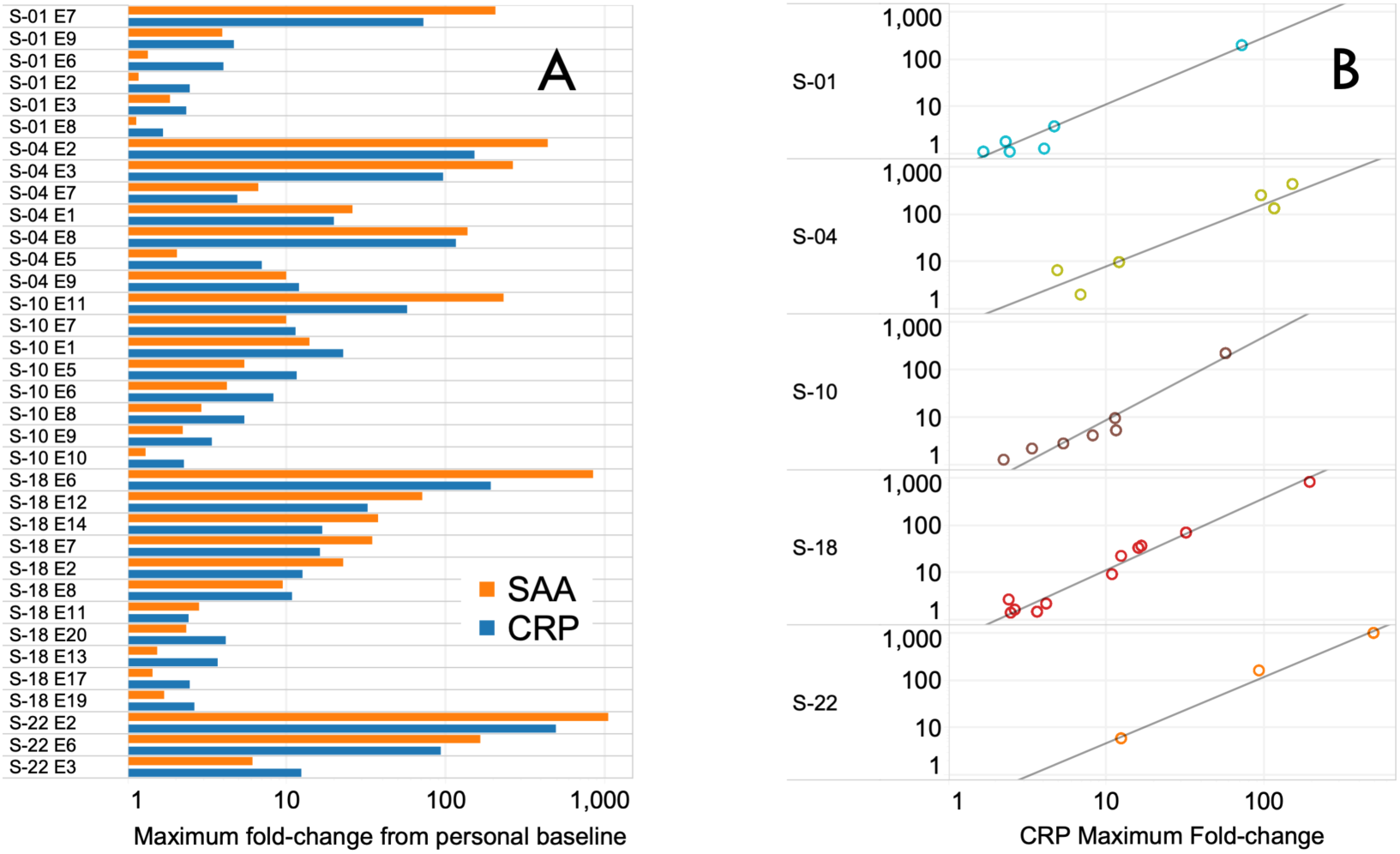
Non-linear relationships between CRP and SAA in response to inflammation. A) Bargraphs showing the maximum fold-increase (log scale) from baseline for SAA and CRP in a series of inflammation events. B) The same events aggregated for each subject (log-log) and fitted with a power model.

### Influenza vaccination

Two subjects collected daily samples in periods that included a vaccination against influenza (Fluzone™ high dose 2017). Inflammatory responses (Figure 9B; S-18 E20 and S-04 E9) in SAA and CRP were measurable (2.2- and 1.3-fold increases in SAA from respective personal baselines), but were ∼100 x smaller than those associated with major infections. Of the 58 identified inflammation events, 41 involved SAA increases larger than 2.2-fold (i.e., a greater response than either of these 2 vaccination events).

### Crohn’s disease

Subject S-10 has severe Crohn’s disease and has diligently explored dietary and other measures to control symptoms. Over a period of ∼2.5yr, CRP and SAA levels were substantially elevated on numerous occasions (Figure 5). Overall, CRP was above 2 times the median personal baseline value in 48% of the individual’s samples, a significantly higher proportion than exhibited by the other subjects (Table 4, Figure 10). A large majority of these samples were collected on a regular weekly (early samples) or daily (later samples) basis, decreasing the likelihood of biases in timing. Over the period of sample collection, S-10 reported success in significantly reducing the frequency and intensity of gut inflammatory events, and remarkably, this was reflected in the significant downward trend in CRP and SAA, reduced intensity of CRP spikes and a progressive decline (almost 3-fold) in Hp, a slowly-responding acute phase protein (Figure 11).

**Fig 10.**
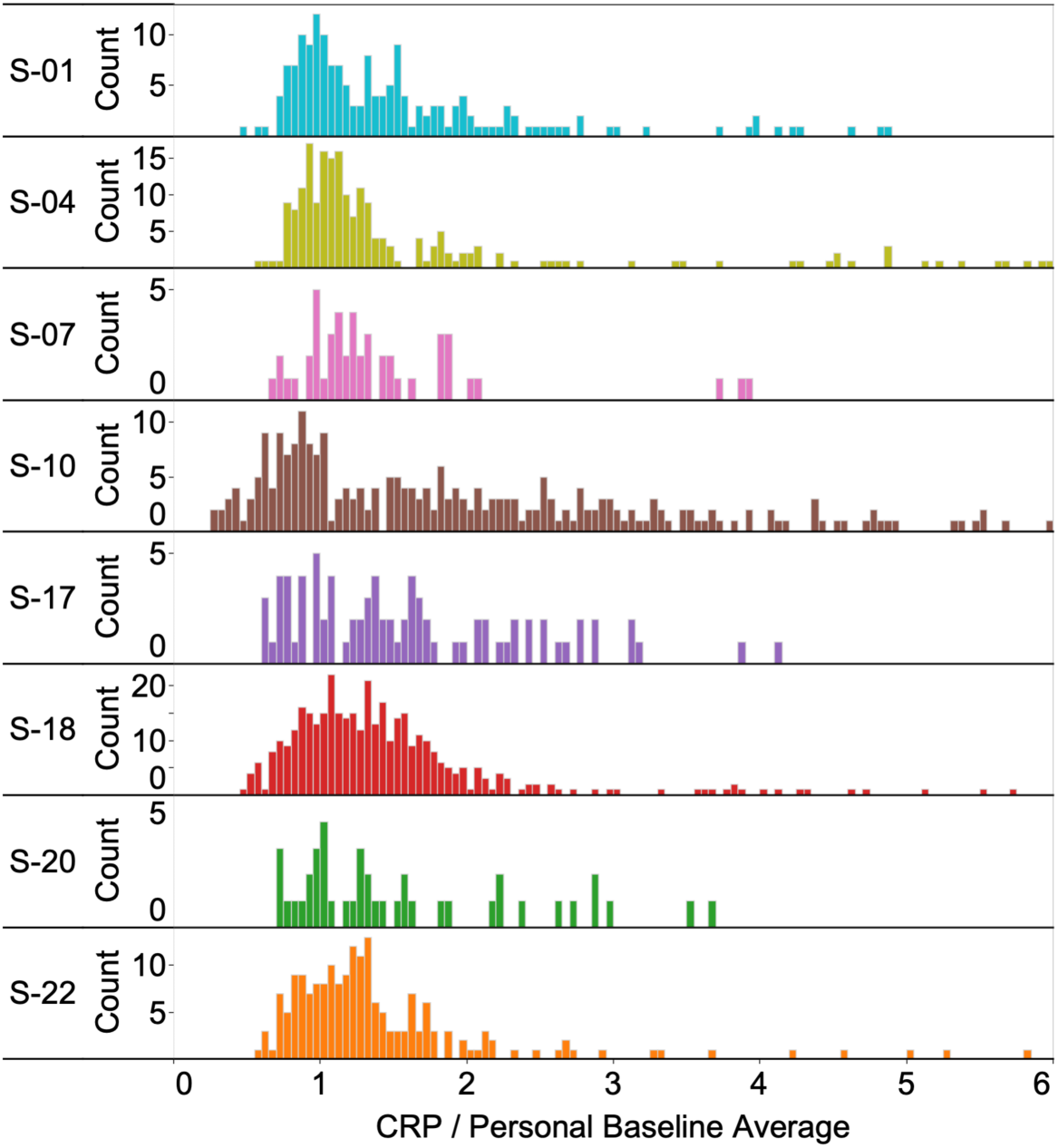
Distribution of personalized CRP values for 8 subjects. Histograms of CRP values (fmol divided by average personal baseline, linear scale focusing on low level variation around baseline) showing greater prevalence of higher levels in Subject S-10 (Crohn’s patient).

**Fig. 11.**
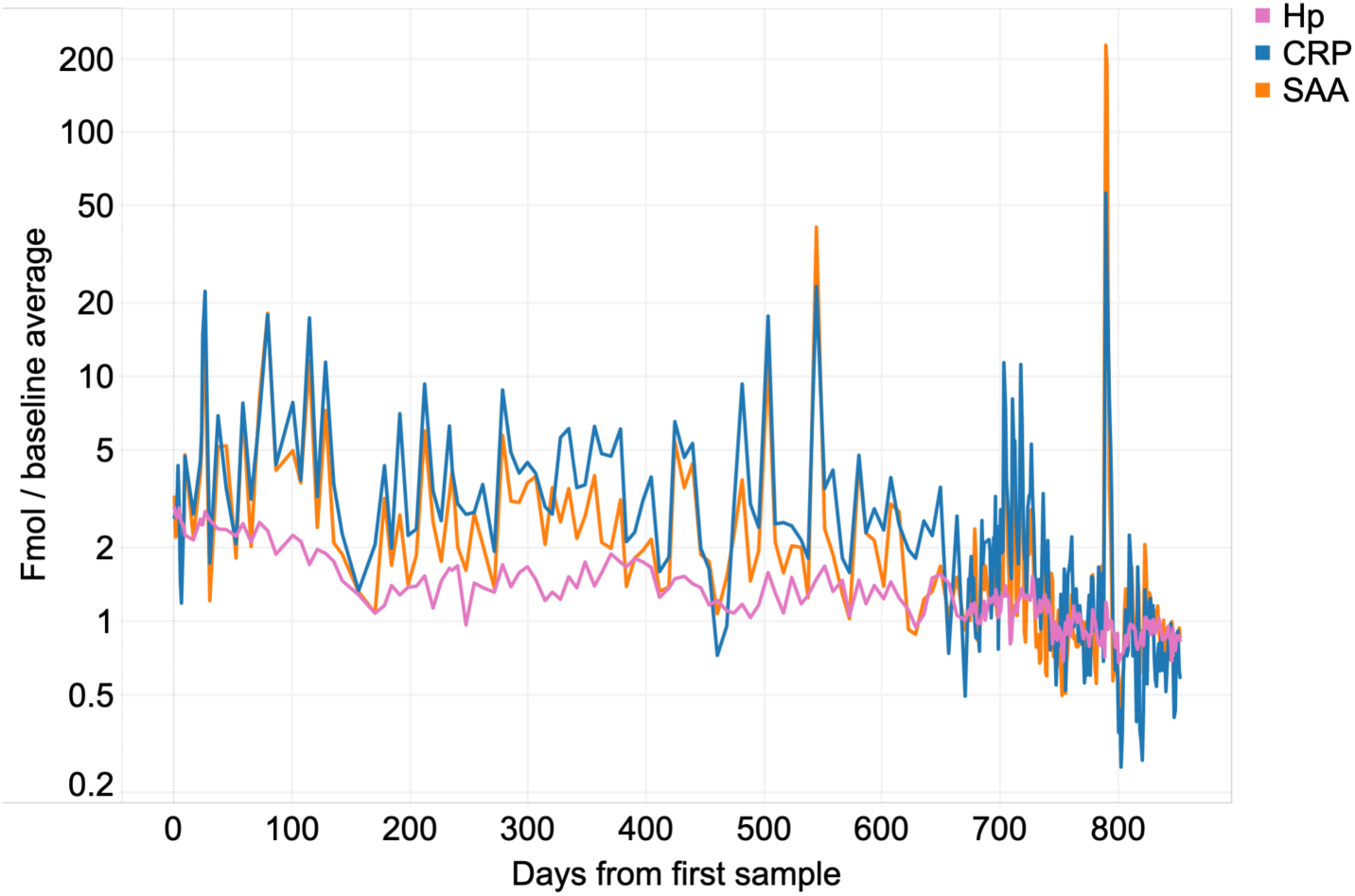
Variation and long-term decline of inflammation markers in subject S-10. Trajectories of declining acute phase markers CRP, SAA and Hp as a patient brings Crohn’s disease symptoms under control. Spike near day 800 corresponds with food poisoning.

### Protein changes secondary to APR protein changes

While the acute phase response showed broad similarity across many infection events in the 8 subjects, episodic responses were observed for MPO and IgM. When measured in whole blood, MPO serves as a surrogate for the neutrophil count (there being much more MPO in the granules of neutrophils present in a volume of blood than in the associated plasma). In 8 inflammation events, increases in MPO preceded the induction of the early inflammation markers such as LPSBP and SAA (Supplementary Figure 5). Instead of a smooth increase and decline shown by the acute phase proteins, MPO exhibited a zig-zag profile with 2-3 days between peaks during persistent infections, perhaps indicative of the hematopoietic cycle time required to produce successive waves of neutrophils in the bone marrow [36].

In contrast, IgM, the dominant early component of the adaptive immune system, showed remarkably stable levels interrupted by significant increases after 3 of the infection events (Supplementary Figure 6) and decreases during most others (where IgM acts as a negative acute phase reactant). In two cases (S-10 E11 and S-18 E6, identified as food poisoning and pneumonia respectively) IgM increased temporarily by ∼50% beginning ∼8 days post-event and then returned to precisely the pre-event level. In a third case, S-01 E7, the increase in IgM was ∼30%. In S-10 E11, where daily samples were available, IgM declined to baseline in ∼15 days, somewhat faster than would be expected based on our measurements of the IgM half-life (∼17d; [37]). No increases in IgG levels (measured as IgGall, the total of all 4 isotypes) were observed.

## Discussion

Precise measurements of protein biomarker panels in self-collected longitudinal DBS samples enables important new opportunities in precision medicine and personalized health monitoring. Here we demonstrate such an approach and describe results obtained using a set of 1,522 DBS samples collected by 8 individuals over intervals of up to 9 years, including extended periods of daily sampling. In these samples we measured panels of up to 27 clinically significant proteins using an automated SISCAPA-LC-MS workflow [5] and normalized the results to personal baselines. As a first step in understanding the major sources of longitudinal biological variation in the human blood proteome, we characterized the strongest driver of change in normal individuals: the acute phase response to inflammation. Given the wide range of proteins and timescales (hours to years) involved in this variation, we conclude that conventional biomarker measurement approaches seriously undersample clinically-relevant biology and thus fail to capture important aspects of health, disease and treatment. Having a practical approach for tracking multiple protein biomarkers at daily (or higher) frequency will overcome this limitation, providing dynamic molecular readouts that can bridge the current gap between the static molecular picture provided by genomics and the dynamic but macroscopic data stream generated by wearable devices.

### DBS-SISCAPA-LC-MS measurement platform

In order to establish dried blood microsamples (e.g., DBS) as a useful alternative to conventional phlebotomy and thereby to enable high-frequency longitudinal tracking of blood biomarker panels, a number of issues must be addressed. These include practicality of frequent sample collection; analytical performance limitations (sample stability, amount of sample required to measure multiple targets, availability of relevant assay panels and overall cost of measurement); volume normalization; and standardization/personalization of resulting data.

DBS samples of fingerprick blood can be collected by most motivated individuals with much less inconvenience [38], cost [39] and risk than conventional venipuncture (venipuncture being considered impractical at daily frequency over extended periods outside a hospital environment and generally discouraged for ethical reasons, such as associated reduction of subject hematocrit [40]). Diabetic patients, for example, can collect 1-5 µL of capillary blood by fingerprick several times each day for glucose measurement and our experience indicates that with proper choice of lancets and technique, daily collection of 50-150 µL blood from a single fingerprick is tolerable for many subjects without injury or undue discomfort. Advanced blood microsample collection devices (e.g., using application of gentle vacuum to collect capillary blood from the upper arm; www.7sbio.com and www.tassoinc.com) will continue to improve user acceptance and likely drive improvements in standardization compared to the first-generation sample collection approach used in this work (handheld lancets and Whatman 903 filter paper cards). Fortunately, the act of drying whole blood samples stabilizes proteins (though perhaps not all small molecules) sufficiently to enable long-term storage at 4° C (as shown in the present results by stable baselines in samples collected over 9 years) and thus enormously simplifies storage of large sample sets. In addition, because the SISCAPA workflow is a “bottom up” proteomics approach in which the target proteins are digested to peptides prior to analysis, the results are much less susceptible to protein stability issues that affect immunoassays. Given these factors, the ability to collect longitudinal DBS samples does not, in itself, appear to be a major limiting factor now or in the future.

Protein analysis, on the other hand, has been limited in DBS. While numerous proteins of clinical interest can be measured in DBS samples [41,42], this is most frequently done with research-use immunoassays that measure one protein at a time, with each assay requiring a separate aliquot of sample and entailing a separate additional cost, thus severely constraining measurement of broad protein panels. Not all existing immunoassays work well in reconstituted dried blood since protein epitopes critical to assay performance can be perturbed by drying and storage. In addition, the lack of knowledge of the volume of plasma in each DBS punch interferes with determination of accurate plasma concentrations of biomarkers required to compare samples with high precision. In contrast, mass spectrometry of protein-specific (proteotypic) peptides offers an ideal approach for measuring many proteins in DBS samples [9]: near-absolute structural specificity; facile, non-interfering multiplex measurement of many peptides; and direct analyte detection with true internal standards (a form of classical isotope dilution mass spectrometry frequently used to establish clinical reference methods). A number of studies have applied MRM methods [43–48] to DBS, demonstrating a capability to measure numerous proteins with reasonable precision and limited throughput. When directed to specific well-characterized peptides, a method combining MS detection with specific immuno-affinity peptide enrichment (SISCAPA; [5,8,32]) significantly enhances the sensitivity, throughput and linear dynamic range of the approach and delivers clinical-quality results [14], making it possible to measure broad panels of proteins in small samples with high precision and at low cost. Here we selected proteotypic peptides from 27 protein analytes for which high-affinity monoclonal anti-peptide antibodies are commercially available (www.siscapa.com/catalogassays). The selected targets span an abundance range of ∼1,000,000-fold in whole blood and include proteins of very high abundance (e.g., HbA and Alb) and low abundance (e.g., IGF1 and sTfR). The assays are highly specific and perform equivalently in plasma or whole blood (manuscript in preparation), with the exception of proteins known to be present in the cellular compartment of blood (HbA, L-plastin, and MPO) which are much more abundant in whole blood than plasma and Hp which interacts very strongly with HbA.

### Interpreting DBS Results: Normalization and Personalization

A central goal of longitudinal DBS analysis is the detection of small changes in an individual over time, a capability requiring very high assay precision (e.g., 2-7% CV). Currently, tests using DBS provide adequate results when used for qualitative analysis (e.g., in screening of newborns for inborn errors of metabolism) or semi-quantitative analysis (e.g., HIV viral load). However, most clinical DBS applications do not attain this level of precision. This is due to variations in the quantity of blood contained in a DBS punch, variation in the fraction of this blood that is plasma (i.e., the hematocrit), differences between capillary and venous blood, and potential alterations in analyte amount or structure due to drying and storage can all affect biomarker measurements. Despite strong interest in DBS technology [49], these concerns have impeded routine application in pharmaceutical research and clinical trials - areas in which detailed longitudinal monitoring of patient responses could be particularly useful [50].

To address these limitations, we have developed [32] and here refined our earlier two-stage strategy to 1) normalize plasma volume and then 2) personalize interpretation of results. In the first step we sought to correct for variations in plasma volume present in each DBS punch – a recognized source of pre-analytic error that typically amounts to +/-10-15% and which adds unacceptable error in longitudinal comparisons. While albumin (Alb) is the largest single protein component of plasma and represents an obvious basis for plasma protein normalization, it is also a negative acute phase reactant and thus its plasma concentration is not always constant. We therefore selected two additional proteins whose concentrations are remarkably stable in individuals over time (IgM and Hx) and which, together with Alb, form a set, in which each protein generally changes by a small amount in compensating directions when major plasma proteome changes occur (e.g., during major inflammation events). To normalize for volume variations in DBS samples, we calculated volume scale factors for Alb, IgM and Hx (for each, the reciprocal of the protein amount divided by the median of the protein amount in all of the subject’s samples) and averaged these scale factors for the three proteins. Consistent with expectation, the distribution of these scale factors across all samples was characterized by a CV of 16.2%, which, if uncorrected, would cause individual assay CVs to substantially exceed this value.

Application of the sample volume scale factor places all of an individual’s samples on an equivalent plasma volume basis, thus normalizing for variation in both the amount of blood in the DBS punch and hematocrit, and resulting in significantly lower assay CV’s. For replicate standards (DBS punches prepared from the same blood sample) run over 9 days in Dataset C, normalization reduced the measurement CV, averaged over 27 peptides, from 10.2% to 4.8%. A number of proteins not used in the normalization showed CVs below 2.5%, close to the limit achievable with current MRM mass spectrometry for peptides. These CVs are generally much lower than most published work with DBS, in which CVs for individual protein assays typically range from 7-20% [41,44,45,47,51]. We attribute this improvement in precision mainly to the combination of highly specific immuno-affinity MS (which effectively separates the peptides analytes from digest matrix) and volume normalization (which has not been available in previous studies where appropriate normalizing proteins were not measured). Lower CVs enable detection of smaller longitudinal changes, and therefore further improvements (e.g., 1% workflow CVs) could further improve signal-to-noise in a number of big data applications.

Interestingly IgM and Hx, while stable within-subject, vary substantially between subjects (i.e., they have a high index of individuality [34]) and as a result, a simple plot of Hx *vs* IgM almost completely separates all of each subject’s samples from the others. This is important evidence, which can be further improved using additional proteins, that a given sample set derives from a single individual despite a span of years in collection times, and provides an excellent tool for detecting mislabeled specimens. In the present study this result also offers a useful example of the long-term stability of these proteins in DBS at 4° C.

Two significant limitations of this plasma volume normalization approach should be noted. First, it is most effective within an individual; i.e., for adjusting samples taken over time from a single individual. This is because the proteins used here for normalization include Hx and IgM, which vary a great deal between subjects (while remaining very stable within an individual). As a result, the method has limited utility for improving comparisons between individuals, as required for cohort studies in a population. A second limitation concerns normalization of proteins associated with non-plasma blood components; e.g., erythrocytes (represented here by HbA) and leukocytes (represented by L-plastin and MPO). This is because plasma volume normalization does not directly address normalization of the whole blood volume, which is affected by blood hematocrit, or potential variations in the ratio of plasma to cellular components due to chromatography during spread of blood across the paper, differential transport during drying, etc.. We address these and other potential sources of small variations in plasma-to-cellular ratios in a separate publication (in preparation). However, neither of these limitations proves to be a disadvantage in the personalized approach we advocate here, since it is based on within-person tracking of plasma protein biomarkers.

As a second step in our analysis we personalized protein amounts in terms of fold-change relative to individual baseline levels, allowing each subject to serve as their own control. This approach has two important advantages. First, it facilitates detection of small changes that are significant in the context of the individual, but may not appear significant in the context of much broader population variation. In the current laboratory diagnostic paradigm, most test results are evaluated against a population reference interval (the normal range defined by results from a population of theoretically healthy people after removing the top and bottom 2.5% of values) - a range that is known to be much wider than an individual’s normal variation for a majority of clinically interesting proteins [52]. Changes in the concentration of a biomarker that are within the population’s “normal” range can be very significant in the individual (i.e., far outside the individual’s narrower personal normal range). The importance of this distinction has been demonstrated by longitudinal monitoring of cancer antigen 125 (CA125) as a predictive biomarker for ovarian cancer [53,54]: individual women exhibit very stable normal levels of CA125 over time but these levels vary widely between women. In such a case, dramatic departures from an individual’s narrow personal range may still be within a much wider population “normal range”, preventing early detection of cancer. While the personalized baseline approach is natural in the context of carefully collected longitudinal biomarker data, it remains very challenging in the context of many existing medical systems due to lack of necessary prior test results and the considerable burden involved in moving from a one-size-fits-all reference interval interpretation to a “complicated” personalized approach. A much more limited version of this personalization concept (use of a Reference Change Value, or RCV, to estimate the significance of differences between sequential measurements) has been proposed [55] but not yet widely adopted.

A second advantage of interpreting biomarker levels relative to personal baselines is that it does not require absolute calibration of each assay in terms of physical concentration (e.g., ng/mL or nM). Absolute physical calibration is highly desirable from a metrological point of view, since, if it can be implemented, it would allow harmonization of results from different laboratories using different detection methods and different vendor platforms. Such universal harmonization has, however, proven very difficult to achieve in practice [56] and is likely to remain so given growing evidence of the genetic and post-translational heterogeneity of many target proteins (leading to different assay responses to material from different subjects) and of the susceptibility of immunoassays to a range of analytical interferences [57]. Experience with multiplexed immunoassays suggests that the effort and complexity of absolute standardization of N assays carried out in the same sample at the same time is at least N-times as difficult as one assay (since each typically requires different calibrators, stripped matrices, purified analytes, etc.) and potentially much more difficult if one is to avoid using separate calibrators for each assay (e.g., 3N calibrators in the present case, which would occupy 3 x 27 = 81 samples of each 96-well sample plate). Thus while a conventional mass calibration approach is possible for SISCAPA panels (using methods previously demonstrated for a number of individual clinical SISCAPA assays [14,15,58]), a simpler alternative, based on the labeled-peptide internal standards used in SISCAPA and other MS methods, would be highly desirable. Quantitative MS-based assays use measured quantities of a stable isotope labeled version of each peptide (SIS) as internal standards and thus provide readouts directly in fmol of target peptide. This approach was originally, but erroneously, referred to as “absolute quantitation” [59]: in fact subsequent experience has shown repeatedly that these values are not absolutely accurate due to several factors including potential loss or modification of some fraction of each SIS peptide after initial quantitation by amino acid analysis (e.g., during storage and processing [60]), and the difficulty of achieving and proving 100% yield even in extremely reproducible tryptic digestion protocols [15]. For these reasons, internally-standardized MS assays can be very precise across a batch of samples where the same SIS cocktail and other reagents are used, but may not remain perfectly accurate in absolute molar terms from batch to batch (where the composition of the SIS cocktail could vary). An effective solution to achieve long-term comparability is use of single point calibrators (aliquots of well-characterized standard samples, as proposed by Hoofnagle [61]) as external standards in each sample batch, or, equivalently, to repeat analysis of a subset of subject samples in different batches, as performed here. Use of a single calibrator is made possible by the wide linear dynamic range of stable-isotope dilution MS detection, as opposed to the sigmoid response of most immunoassays. When comparisons between individuals, or with population values, is the objective, or when comparability with existing clinical assay methods is required, absolute values can be assigned to a calibrator [61] to link results to established reference levels. Using the single point calibrator approach, values can be assigned to the standard samples at any time, including the future, and the results propagated backwards to retroactively calibrate previous analyses.

### Scale and Scope of Inflammation Responses

As a first example of high-frequency longitudinal DBS analysis we explored the most common source of multivariate change in the plasma proteome: inflammation. Episodic acute phase responses to inflammation represent the largest effects observed in this data set, with numerous examples spanning a broad range of intensities, involving many proteins and driven by a variety of causes. Previous studies have measured protein panels by a variety of methods to track inflammation associated with infection [62,63], typhoid vaccination [64], surgery [12] and LPS-administration [65] at a small number of time points (e.g., 2-7), but none have had the assay precision, time resolution and breadth of causation available in the present data set. Among 8 subjects in this pilot study, all of whom (except for the single Crohn’s patient) were generally healthy, we observed inflammatory events related to: surgery (a short duration inflammatory pulse followed by a smooth recovery); infections (following variable courses ranging from days to weeks, each having a discrete beginning and end); and Crohn’s disease (a chronic pattern of recurring inflammation with components lasting years).

The largest quantitative changes we observed in each subject resulted from infection, and included increases of more than 1,000-fold in SAA and more than 100-fold in CRP, accompanied by smaller increases in proteins such as LPSBP, MBL, A1AG, Hp, FibG, Hx and C3 (typically in a range of 1.2-to 4-fold). A less dramatic event (influenza vaccination) drove SAA increases of only 1.3-to 2.2-fold (consistent with published studies [66]) – yielding a dynamic range of almost 1000 in detectable APR responses. Even smaller amplitude “microinflammation events” were detectable through the coordinated behavior of SAA, CRP and LPSBP. These inflammation biomarkers showed significant correlations (Figure 3) even in the half of each subjects’ samples with lowest CRP, demonstrating that low-level variations are not due simply to measurement noise but instead contain measurable inflammatory modulations hidden in the baseline values.

Each APR protein followed a characteristic time course, most clearly visible in the response to a short surgical procedure (a 2 hr total hip arthroplasty). Because of these differences and despite substantial correlations observed among the markers, all of which are produced primarily in the liver in response to cytokine signaling [19], a set of APR proteins provides much more information over time than CRP alone. For the positive APR, the time order of increase is typically LPSBP, SAA, CRP, A1AG, FibG, Hx, MBL, Hp, C3, with the more rapidly increasing proteins (SAA, CRP, LPSBP) showing the greatest quantitative changes relative to baseline. The order of decline is similar but not identical (SAA, CRP, LPSBP, Hp, Hx, FibG, MBL, A1AG, C3), an order that is generally consistent with previously measured half-lives of CRP, LPSBP, A1AG and C3 of 26, 32, 104 and 84 hr respectively [37]. APR proteins also showed complex amplitude control. Peak levels of SAA and LPSBP achieved in inflammation events are non-linearly related to peak CRP. Indeed, SAA shows a greater fold-increase over baseline than CRP in stronger APR responses, while the opposite is true in responses to low level infections. These results suggest that a dimensionless ratio of SAA and CRP fold-change relative to personal baseline could be a useful indicator of infection severity.

As a first step to move beyond conventional single-parameter CRP measurements of inflammation we used simple 2-parameter plots [35] to track the sequence of events involved in departing from, and then returning to an individual’s healthy baseline state of inflammation. An example (Figure 7) plotting SAA (here summarizing the rapid component of the APR response) *vs* Hp (slower APR response) depicts recovery following hip replacement surgery as a counterclockwise loop, tracking inflammatory physiology independent of clock time. This smooth recovery loop is interrupted by renewed inflammation at several points, coinciding with the subject’s reports of a localized pelvic strain and resumption of pain medication. A 50% decline in inflammation level between un-medicated pre-operative and post-surgical baseline samples provides a quantitative demonstration of the reduction in inflammation following replacement of a diseased hip joint. Both effects suggest the potential value of inflammation markers as surrogates for pain in the context of tissue damage. Similar loop presentations of a series of infection events (Figure 8) indicate differences in magnitude and path that likely relate to the etiology and severity of infections (though no infectious agents were identified in these cases as none required hospitalization).

Taken together, the complex time and amplitude relationships among the APR proteins suggest that advanced multiparameter modeling approaches like those used in pharmacodynamics [67] and machine learning [2] will emerge as valuable tools for inflammation monitoring. Such models can be personalized based on past infection loop histories (e.g., training the models on past colds or vaccinations to “calibrate” a subject’s responses) and used to improve assessment of both infection severity and stage of response (i.e., where the subject is in the cycle of infection and recovery, independent of clock time).

A surprisingly high proportion (12-45%) of each subject’s samples showed indications of inflammation, whether defined as CRP levels >2-fold above personal baselines, or inclusion in the 58 discernable APR events. Similar proportions are obtained when considering only those samples collected on a near-daily basis (about 1/3 of the samples), in which the timing of collection should be less susceptible to subjects’ selection bias. Such high frequencies, if confirmed in a larger subject population, suggest that unrecognized short-term inflammatory events are likely to occur during well-designed investigational studies including drug trials. Numerous small molecule drugs are known to bind to albumin and/or A1AG [68], both of which are observed to change significantly in inflammatory situations, potentially altering drug distribution in both trials and clinical use. Likewise randomly-timed single CRP measurements (such as those included in annual checkups) may frequently overestimate a patient’s true baseline level if collected during a microinflammatory episode, and thus bias predictions of cardiovascular disease risk [18,69]. The use of a personal baseline CRP level derived from a series of longitudinal samples should provide a better risk estimate by capturing baseline inflammation separately from fluctuations related to transitory subclinical events.

In addition to acute phase proteins, we also observed increases in IgM and MPO in a subset of infections. While most infections did not cause measurable increases in IgM, three episodes resulted in increases of 30-50% in total IgM 8-10 days after SAA peaked, after which the levels subsided to pre-existing baseline levels. The production of such large amounts of IgM after specific infection events is consistent with the expected timeframe for an adaptive immune response to a pathogen, and may provide an opportunity to identify endogenous human monoclonal antibodies from DBS samples with potential therapeutic value as anti-infectives. In contrast, MPO, which serves in whole blood as a surrogate for neutrophil count, was frequently increased at the earliest stage of infection followed by spikes every 2 or 3 days, consistent with periodic release from and regeneration of neutrophil pools in the bone marrow [36].

### Future outlook for dense longitudinal analysis

Longitudinal measurements of clinically established blood protein biomarkers can be considered as dynamic readouts from internal physiological sensors and as such it is natural to envision combining them with other emerging dynamical health data sources [70] including heart-rate, steps, sleep and electrocardiogram data from smart watches and phones; data collected by continuous positive airway pressure (CPAP) devices used to treat apnea; glucose sensors; local weather records; as well as a wide variety of user-provided “contextual data” collected though disease-centric (e.g., Crohn’s disease), weigh-loss and dietary smartphone apps. Combinations of protein biomarkers with these data streams will play an increasing role in data interpretation as we report additional results on other components of the DBS panel measured here related to lipoproteins, iron metabolism, kidney function, etc.. Taken together, dynamic measurements from internal physiological sensors (blood proteins) and external mechanic-physical sensors will provide “Big Data” of sufficient size, scope and precision to enable productive machine learning and the construction of practical personalized health models.

## Materials and Methods

### Samples

Capillary blood samples (1,522 in total, Table 1) were self-collected by participants at home using lancet finger-pricks (Medlance Plus Extra or Special, HTL Strefa) and dried on Whatman 903 Proteinsaver dried blood spot (DBS) cards (5 drops on each card). Participants provided informed consent under IRB approval (Quorum Review 33239CDN/1). The DBS cards were generally stored at 4° C in the presence of desiccant except for brief periods at room temperature or at −20° C, and were barcoded prior to analysis. Transportation of specimens to the laboratory for processing followed the guidelines provided by the Center for Disease Control and Prevention (CDC) for shipment of DBS specimens. Individual samples consisted of punches taken from blood-covered regions of DBS cards and deposited in wells of 96-well plates (four 1/16” punches for Dataset A and a single ¼” punch for Datasets B and C).

### SISCAPA-LC-MRM protein measurement

Groups of samples were analyzed on three separate occasions (in 2015, 2016 and 2017) to generate datasets identified here as Datasets A, B and C. Sample preparation and SISCAPA enrichment were performed using an automated protocol essentially as described [5] for all three datasets, with different, but largely overlapping, sets of measured protein targets (TABLE 2). In this protocol, proteotypic tryptic peptides (whose sequence appears only in the target protein) were used as surrogates for proteins after digestion of DBS samples with trypsin (one peptide per target protein, except for Apo A-I and Apo B lipoproteins when two were used for each). These peptides were quantitated in relation to added same-sequence stable isotope labeled peptide internal standards (SIS) by MRM mass spectrometry after immuno-affinity enrichment using anti-peptide antibodies (SISCAPA) as previously described [5,8,32,71]. The protocol is implemented as a series of liquid addition steps to dissolve and denature DBS proteins, followed by disulfide reduction and alkylation, tryptic digestion, addition of stable isotope labeled (“SIS”) versions of the target peptides as internal standards, capture of target tryptic peptides and internal standards by anti-peptide antibodies (one for each target peptide) on magnetic beads, and finally washing of the beads and elution of target peptides for injection into the LC-MS, which then measures the peak area ratio of endogenous sample-derived tryptic peptides to their corresponding stable-isotope labeled internal standards. This automated “addition only” protocol is designed to increase precision [5]. Target protein amounts were expressed as femtomole (fmol) of proteotypic peptide in each sample, calculated by multiplying the observed MRM peak-area ratio (endogenous target : SIS) by the known amount of added SIS. SIS peptides were labeled with 13C, 15N lysine or arginine as the c-terminal amino acid, except in the case of ApoE where the proteotypic peptide was at the c-terminus of the protein and an internal valine residue was labeled instead.

Samples were analyzed in three tranches on separate occasions: Datasets A, B and C, run in 2015, 2016 and 2017 covering 20, 12 and 27 peptides in 600, 423 and 606 samples respectively (TABLE 2). Dataset A results (792 samples from 16 subjects) have been previously described in part [32], and used an LC-MS/MS platform consisting of an Agilent 1290 LC operating at 600 μL/min and an Agilent 6490 triple quadrupole MS. Replicate standards were prepared as punches of whole blood dried on 903 paper (4 punches of 1.5 mm diameter were used per well).

Datasets B and C employed an Eksigent microflow LC system operating at 10 μL/min connected to a Sciex 6500 Q-trap MS. Dataset B (436 samples from 4 subjects) focused on measurements of a panel of gastrointestinal inflammation related proteins and employed a single MRM for each peptide. Replicate standards were prepared by pipetting whole blood onto pre-cut 6 mm diameter disks of 903 paper in sample wells.

Dataset C (792 samples from 10 subjects) was comprised of two sequential panels applied to the same DBS digests and employed 1 to 3 MRMs for each peptide (PAR for the selected MRMs were averaged to generate a single value for each peptide measured in a sample). All the panels and plexes included the normalizing protein triad (albumin, hemopexin and IgM). Replicate standards were prepared as punches of whole blood dried on 903 paper.

Nine peptides were measured in all datasets (including Alb, Hx, and IgM) and 19 were measured in both Datasets A and C. As part of continuing efforts to increase assay precision, the 27-plex used in Dataset C was measured as two sequential SISCAPA multiplex subpanels, both of which included normalizing proteins Albumin (Alb), Hemopexin (Hx) and total immunoglobulin M (IgM). Datasets A and B used a single MRM per peptide (measured as a single multiplex panel), while Dataset C used the average of 1 to 3 of the best quality MRM transitions available for the respective peptides. Datasets B and C were each merged with Dataset A using a run-to-run scale factor for each protein in common between datasets in order to account for potential shifts in the absolute amounts of the SIS peptides during storage between the analytical runs. The scale factor was derived from the ratio of average values for a protein in subject samples that were analyzed in both Datasets (88 samples analyzed in both Datasets A and B, and 19 samples analyzed in both Datasets B and C, in each case using duplicate punches from the same DBS card). When samples from the same subject were shared between datasets this ratio was subject-specific; for subjects without shared samples an average of the subject-specific ratios was used.

### Data analysis

Peak area ratio (PAR) measurements in Dataset A were obtained using MassHunter Quantitative Analysis (v. B.05.02), while in Datasets B and C the Sciex MultiQuant program (v. 3.0.2) was used. PAR data was assembled in Tableau Prep Builder (www.tableau.com) and joined with tables defining sample characteristics (e.g., date of collection, subject contextual health notes, etc.), SIS concentrations, analytical run structures, etc.. Data analysis and visualization was carried out in Tableau.

## Acknowledgments

We thank Jordan Kallman and Steven Skates for helpful, insightful discussions, and all sample donors.

## Funding

SISCAPA Assay Technologies, Inc.

## Author contributions

NLA and MR designed the project with input from MP and TWP, MR and RY carried out analytical workflow, NLA carried out data analysis in Tableau and wrote a draft manuscript which was edited by MR, MP and TWP.

## Competing interests

All authors are affiliated with SISCAPA Assay Technologies, Inc., which has obtained patents on the SISCAPA sample preparation workflow.

## Data and materials availability

## Supplementary Materials

**Supplementary Table 1.** Impact of volume normalization on coefficients of variation of replicate standards in three datasets.

| Protein | Dataset A Standards |  | Dataset B Standards |  | Dataset C Standards |  |
| --- | --- | --- | --- | --- | --- | --- |
|  | CV Before Vol Normalization | CV After Vol Normalization | CV Before Vol Normalization | CV After Vol Normalization | CV Before Vol Normalization | CV After Vol Normalization |
| A1AG | 11.3% | 8.6% |  |  | 9.2% | 4.7% |
| Alb | 7.2% | 1.9% | 7.2% | 5.7% | 10.4% | 1.9% |
| ApoA1 (AKP) |  |  |  |  | 10.0% | 1.5% |
| ApoA1 (ATE) | 7.9% | 4.6% |  |  | 11.2% | 6.5% |
| ApoB (FPE) | 6.2% | 2.4% |  |  | 9.7% | 2.4% |
| ApoB (TEV) |  |  |  |  | 10.5% | 2.8% |
| ApoC-III |  |  |  |  | 9.4% | 4.2% |
| ApoE | 7.2% | 5.5% |  |  |  |  |
| ATIII | 12.4% | 9.4% |  |  | 8.0% | 9.6% |
| C3 | 6.2% | 2.3% |  |  | 9.3% | 5.5% |
| CRP | 14.1% | 13.1% | 6.8% | 6.8% | 9.8% | 3.3% |
| CysC | 5.6% | 6.1% |  |  | 11.7% | 3.5% |
| FibG | 6.9% | 5.3% |  |  | 10.6% | 4.3% |
| HbA | 7.0% | 5.6% | 7.3% | 7.0% | 10.3% | 4.9% |
| Hp | 10.2% | 9.4% | 5.3% | 5.5% | 6.8% | 9.4% |
| Hx | 6.4% | 1.4% | 4.6% | 3.0% | 10.0% | 1.5% |
| IGF-1 | 20.4% | 19.8% |  |  | 9.8% | 2.0% |
| IgGall |  |  |  |  | 10.6% | 5.8% |
| IgM | 6.1% | 2.6% | 6.1% | 4.6% | 9.8% | 2.1% |
| L-Plastin |  |  |  |  | 11.1% | 5.0% |
| LPSBP | 7.6% | 6.5% | 6.4% | 6.2% | 10.2% | 3.7% |
| MBL | 8.1% | 7.8% | 11.1% | 11.2% | 14.2% | 12.1% |
| MPO | 11.7% | 13.1% | 5.8% | 5.5% | 10.5% | 9.7% |
| Pla | 6.7% | 2.8% |  |  | 9.9% | 5.0% |
| SAA | 10.9% | 8.8% | 5.0% | 4.8% | 11.6% | 3.7% |
| sTfR |  |  |  |  | 9.8% | 7.1% |
| Tf |  |  |  |  | 9.6% | 2.5% |
| TIMP1 | 9.5% | 8.2% |  |  |  |  |
| vWF | 18.1% | 18.2% |  |  |  |  |
| Average | 9.4% | 7.4% | 6.5% | 6.1% | 10.2% | 4.6% |

**Supplementary Table 2.**
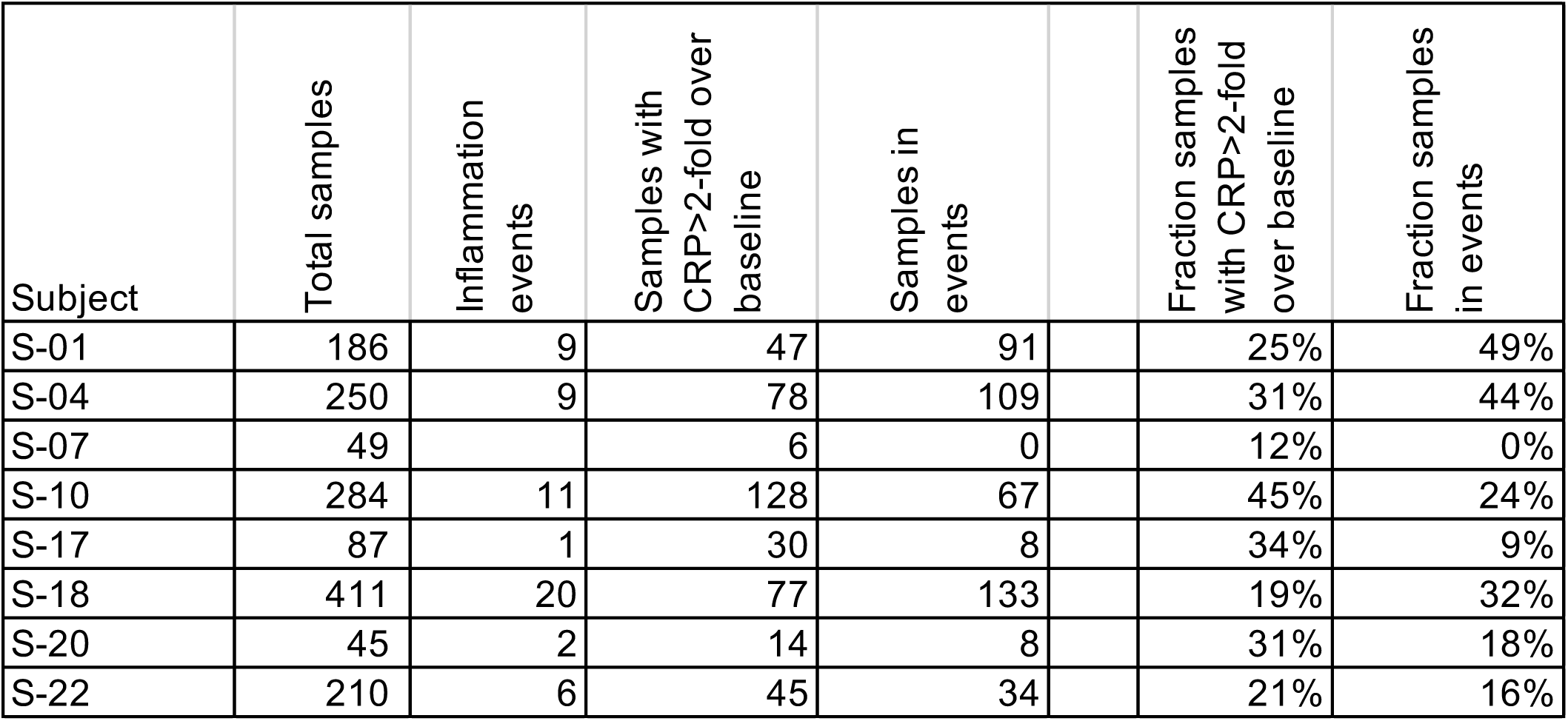
Proportion of subject samples with evidence of inflammation.

**Supplementary Fig 1.**
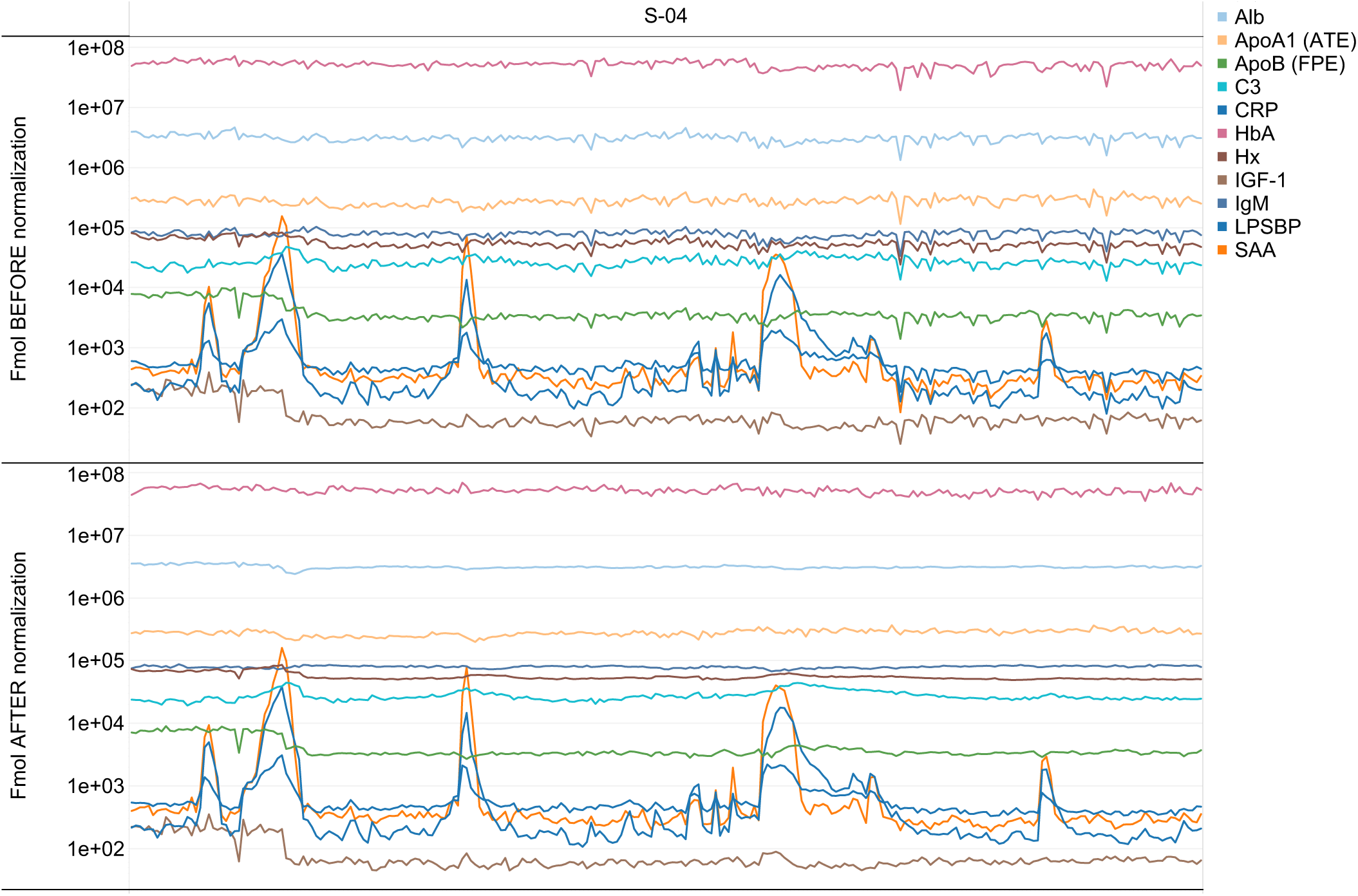
Amounts (fmol of peptide) for 11 proteins measured in 255 longitudinal DBS samples from a single subject before (top panel) or after (lower panel) DBS volume normalization by Alb, Hx, IgM.

**Supplementary Fig 2.**
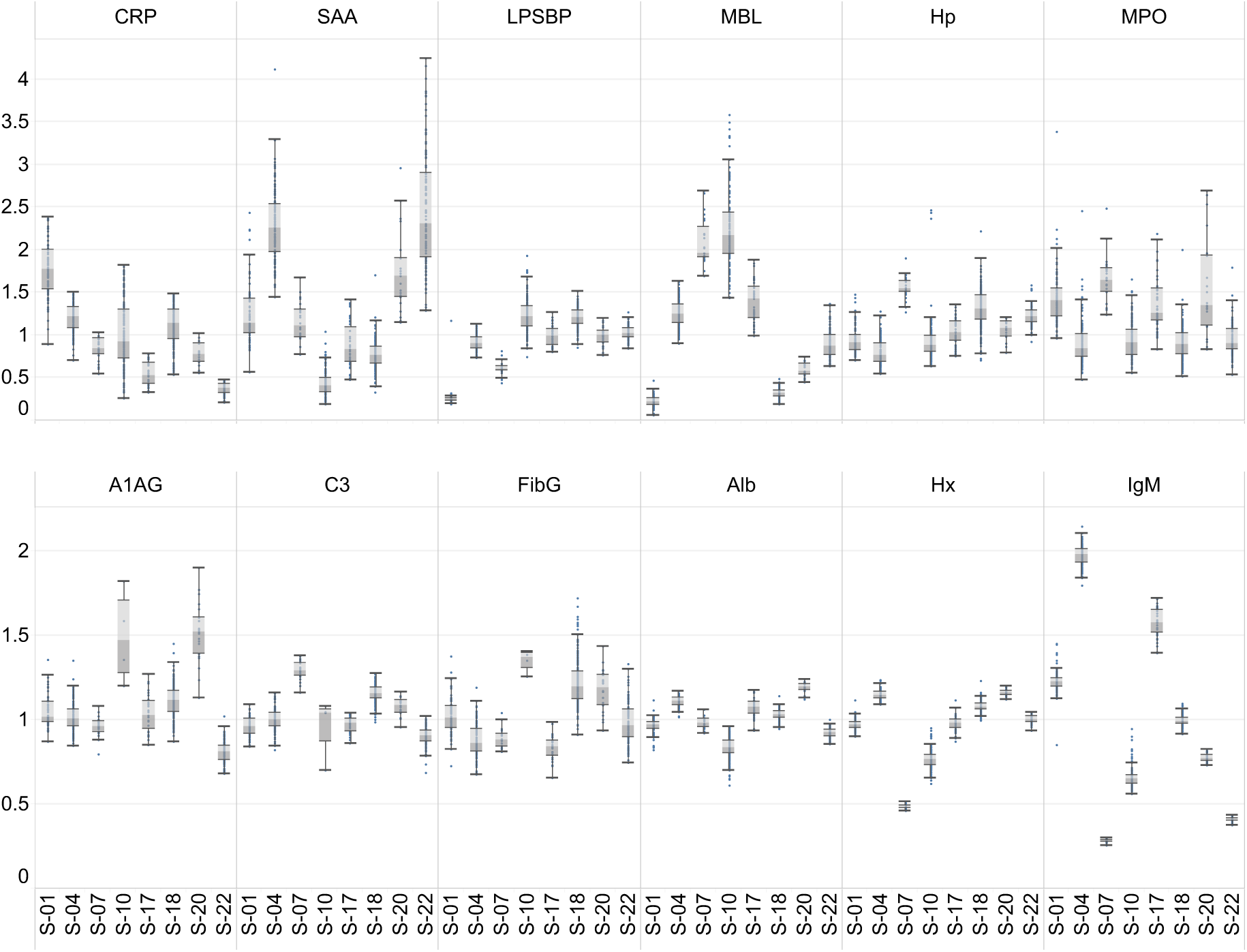
Whiskerplots of the amounts of 12 proteins in the baseline samples of each of 8 subjects. Protein amount has been normalized between proteins by dividing by the median amount in all subjects.

**Supplementary Fig 3.**
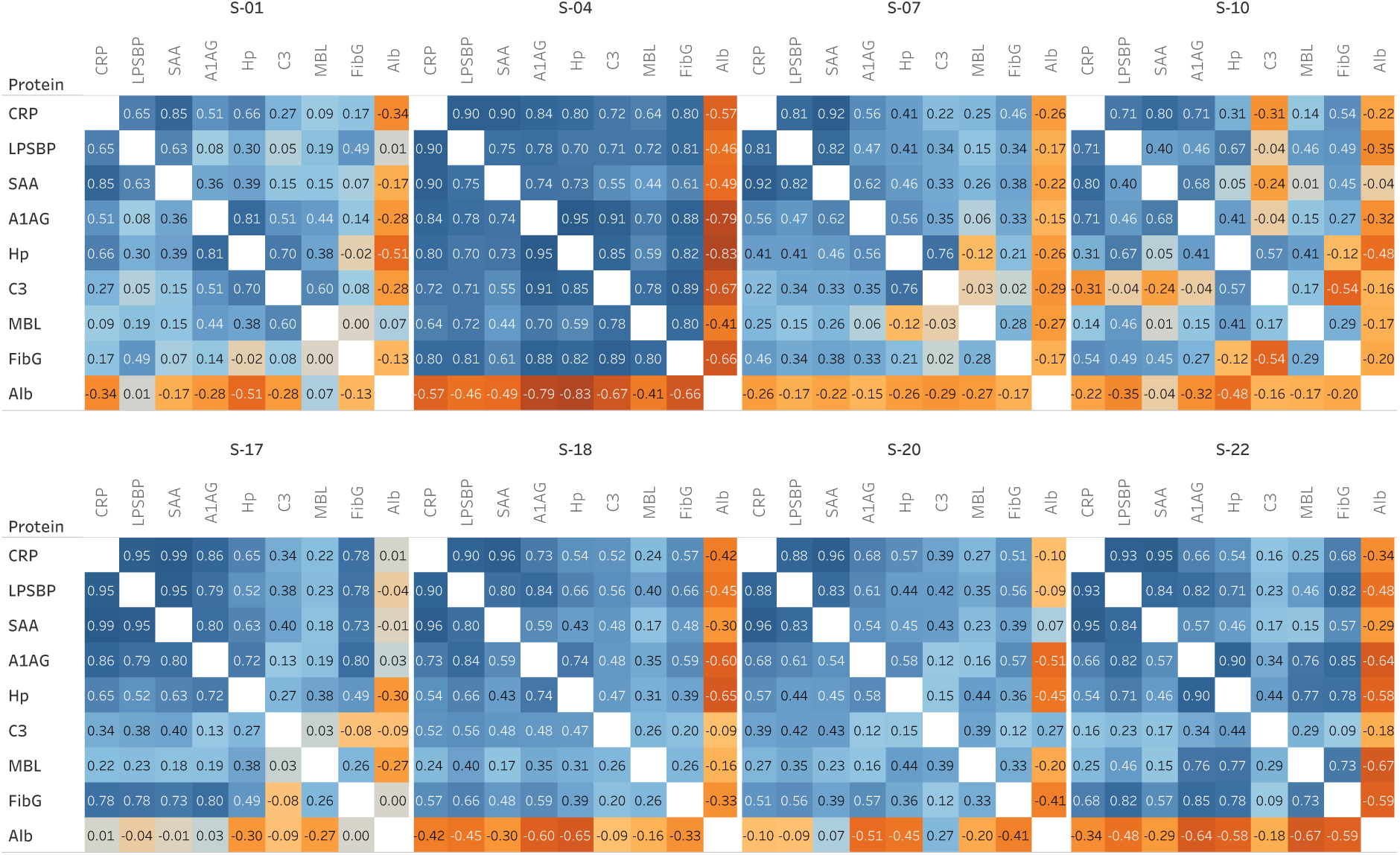
Protein:Protein correlation matrices for inflammation-related proteins calculated using all samples from each subject separately after volume normalization and division by personal baseline average values.

**Supplementary Fig 4.**
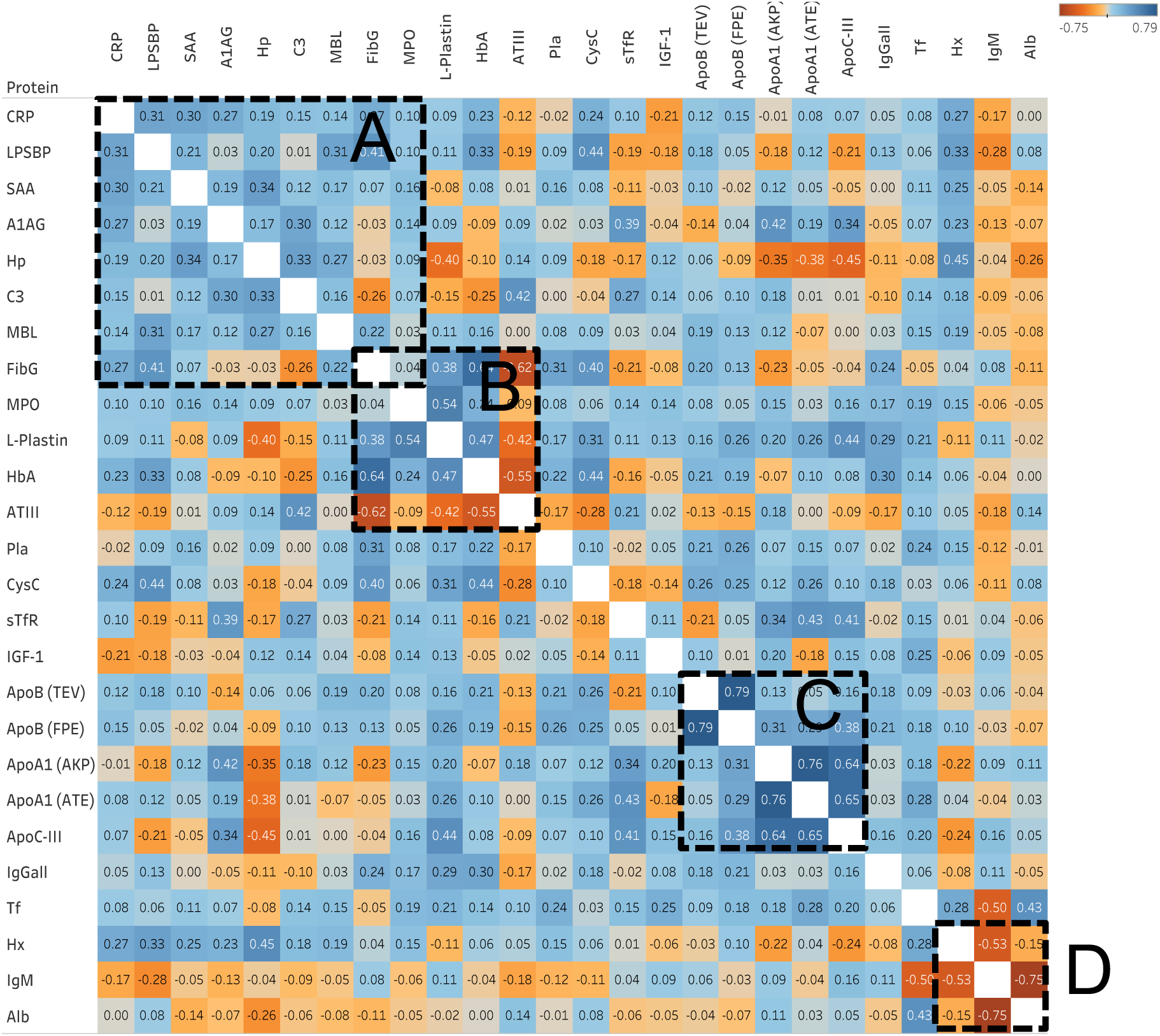
Protein:Protein correlation matrix calculated using only the baseline (non-inflammation) samples from each subject after volume normalization and division by personal baseline average values. A: acute phase response proteins; B: cluster related to coagulation and blood viscosity; C) lipoproteins; D) proteins used in volume normalization.

**Supplementary Fig 5.**
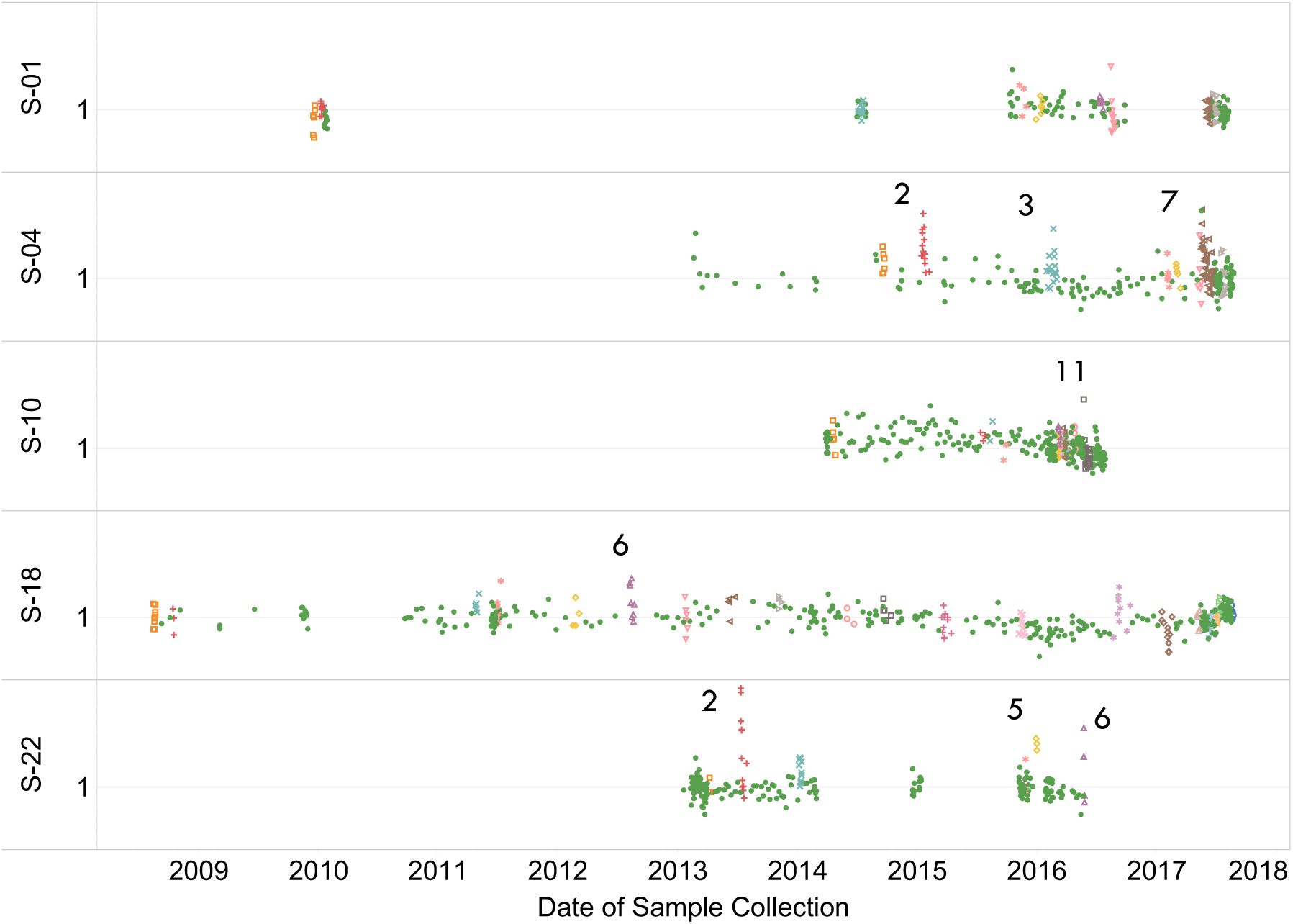
Values for MPO normalized by personal baseline showing several infection events in which MPO temporarily increased.

**Supplementary Fig 6.**
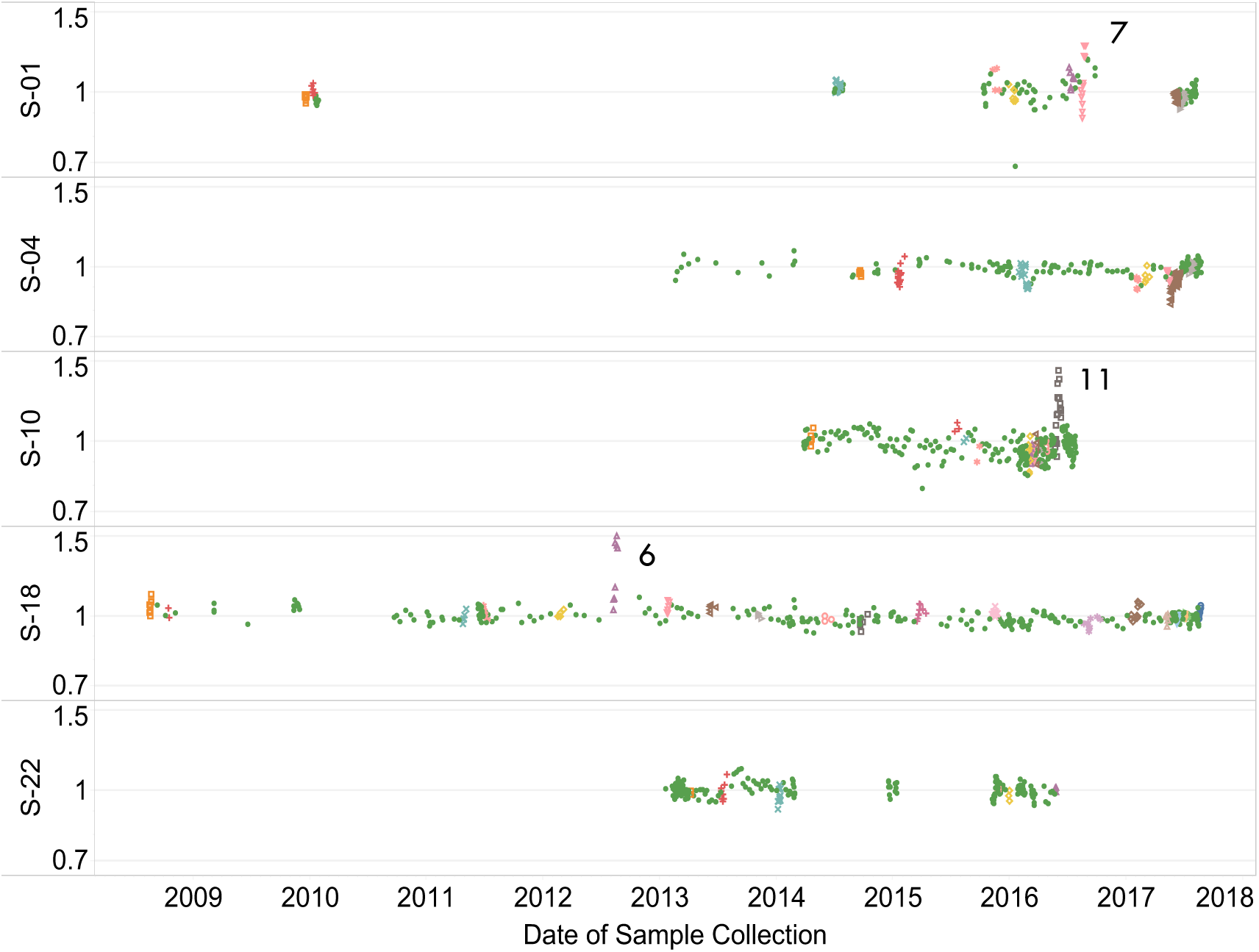
Values for IgM normalized by personal baseline showing two infection events (S-10 E11 and S-18 E6) in which IgM temporarily increased by more than 50% and the returned to baseline levels. A third event (S-01 E7) showed a temporary increase of about 30%.

